# Human embryos arrest in a quiescent-like state characterized by metabolic and zygotic genome activation problems

**DOI:** 10.1101/2021.12.19.473390

**Authors:** Yang Yang, Liyang Shi, Xiuling Fu, Gang Ma, Zhongzhou Yang, Yuhao Li, Yibin Zhou, Lihua Yuan, Ye Xia, Xiufang Zhong, Ping Yin, Li Sun, Zhang Wuwen, Isaac A. Babarinde, Yongjun Wang, Xiaoyang Zhao, Andrew P. Hutchins, Guoqing Tong

**Affiliations:** Center for Reproductive Medicine, Shuguang Hospital Affiliated to Shanghai University of Traditional Chinese Medicine, Shanghai, 200120, China; Shenzhen Key Laboratory of Gene Regulation and Systems Biology, Department of Biology, School of Life Sciences, Southern University of Science and Technology, Shenzhen, 518055, China; BGI Genomics, BGI-Shenzhen, Shenzhen, China; Longhua Hospital, Shanghai University of Traditional Chinese Medicine, Shanghai 200032, China; Key Laboratory of Theory and Therapy of Muscles and Bones, Ministry of Education, Shanghai 200032, China; State Key Laboratory of Organ Failure Research, Department of Developmental Biology, School of Basic Medical Sciences, Southern Medical University, Guangzhou, Guangdong, 510515, China; Guangdong Key Laboratory of Construction and Detection in Tissue Engineering, Southern Medical University, Guangzhou, Guangdong, 510515, China; Guangzhou Regenerative Medicine and Health Guangdong Laboratory (GRMH-GDL), Guangzhou, Guangdong, 510700, China

**Keywords:** Quiescence, Embryonic development, metabolism

## Abstract

Around 60% of *in vitro* fertilized (IVF) human embryos irreversibly arrest before compaction between the 3-8-cell stage, posing a significant clinical problem. The mechanisms behind this arrest are unclear. Here, we show that the arrested embryos enter a quiescent-like state, marked by cell cycle arrest, the downregulation of ribosomes and histones and downregulation of MYC and p53 activity. Mechanistically, the arrested embryos can be divided into three types. Type I embryos fail to complete the maternal-zygotic transition, and type II/III embryos have erroneously low levels of glycolysis and variable levels of oxidative phosphorylation. Treatment with resveratrol or nicotinamide riboside (NR) can partially rescue the arrested phenotype. The mechanism of reactivation involves the upregulation of SIRT1, and activation of glycolysis and fatty acid oxidation which forces the embryos out of a quiescent state. Overall, our data reveal how human embryo arrest can be overcome by modulating metabolic pathways.

## Introduction

*In vitro* fertilization (IVF) has revolutionized the treatment of human fertility problems. However, a large number of human embryos fail to develop *in vitro*, and typically only 30% of human embryos will progress to the blastocyst stage (Alikani et al., 2000; Wong et al., 2010). Human pre-implantation embryos can arrest at all stages between the zygote and the blastocyst. However, a large fraction irreversibly arrest between the 2-cell and 8-cell stages and remain un-compacted (Alikani et al., 2000). Several cellular mechanisms have been proposed to explain this arrest, specifically: failed zygotic genome activation (ZGA) (Vera-Rodriguez et al., 2015), delayed maternal RNA clearance (Sha et al., 2020), reactive oxygen species causing endoplasmic reticulum stress (Kawamura et al., 2010; Rocha-Frigoni et al., 2015), and aneuploidy (Almeida and Bolton, 1998). Computational machine learning techniques can detect morphological patterns in microscope images of otherwise normal-appearing embryos that will later go on to arrest (Wong et al., 2010), suggesting the arrest mechanisms are active before they manifest. However, the cellular mechanism remains unclear.

*In vitro* developmental models of embryogenesis in other organisms has not brought clarity to this problem, as pre-implantation development is divergent between species (Liu et al., 2021a). For example, ZGA mainly occurs at the 2-cell stage in mice, but in humans, there are two waves, a minor ZGA at the 2-cell stage and the major ZGA at the 8-cell stage (Schulz and Harrison, 2019). Relatedly, in contrast to humans, some species have good *in vitro* developmental potential. For example, ∼90% of mouse, ∼80% of (monospermic) pig, ∼70% of cat, and ∼60% of *Macaca mulatta* embryos successfully develop to the blastocyst stage (Gil et al., 2010; Herrick et al., 2007; VandeVoort et al., 2009). Conversely, humans are not the only species with poor *in vitro* embryonic developmental potential, only 25-30% of cattle and horse embryos will develop to a blastocyst (Dinnyes et al., 1999; Hinrichs, 2010; Rocha-Frigoni et al., 2015). However, it is unclear if the same mechanisms are active in other species. Human embryonic stem cells can be manipulated to form artificial blastocyst-like ‘blastoids’ which mimic natural blastocysts (Liu et al., 2021b; Yu et al., 2021). These might be used to understand embryonic arrest. Interestingly, blastoids are generated at low efficiency, which may reflect developmental problems inherent to natural blastocysts. However, blastoids cannot address pre-morula developmental arrest, as they model a later developmental stage, and it is unclear if the problems seen in blastoids are the same as pre-implantation embryos. Ultimately, to investigate the arrest of human embryos, it is necessary to assay the problems directly.

In this study, we utilized genomic technologies to explore the transcriptomic basis behind the arrest of human embryos. A subset of the arrested embryos enter into a quiescent-like state characterized by the up-regulation of p53, MYC, FOXO1, and the widespread down-regulation of ribosomes, histones and translation initiation factors. We show that this quiescent phenotype can be partially reversed using the antioxidant resveratrol, and nicotinamide riboside (NR), and our data suggest that these two molecules activate the sirtuin family of acetyltransferases (SIRTs) to modulate metabolism. Modulation of SIRT activity leads to a reactivation of the arrested embryos, and progression to a morula and early blastocyst. We show that resveratrol and NR alter the balance of metabolism, and the arrested embryos activate glycolysis and fatty acid-oxidation metabolic pathways. Ultimately, these observations can explain at least some features of the arrest of human embryos.

## Results

### Gene expression of arrested human embryos after *in vitro* fertilization

Under typical IVF procedures, ∼60% of embryos arrest (**Fig. 1A**). We were interested in the class of embryos that arrest during development, but maintained a normal morphology and cell integrity, and did not show signs of disintegration. Embryos often arrest at either day 3 or day 4, post-fertilization (**Fig. 1A**). Day 4-arrested embryos reach the 8-cell stage but would fail to form a morula. The day 3-arrested embryos would undergo cleavage, but failed to reach the 8-cell stage (**Fig. 1B, C**). In this study, we focused on the day 3-arrested embryos. The embryos were left a further day to confirm no further development and no fragmentation or disintegration of the embryonic cells. Under these criteria, the embryos would be discarded in IVF procedures. It should be noted that, by this definition, ∼3% of the day 3-arrested embryos can spontaneously recommence development and form a blastocyst (see later in the manuscript). However, prolonged *in vitro* culture of human embryos is deleterious for further development (Alikani et al., 2000), hence we chose the minimal window between confirming the arrest of the embryos and preserving developmental competency. For this study, we defined irreversibly-arrested as day 3 embryos at the 2-cell to 5-cell-stage that remained in that state on day 4, at which point a normal human embryo would be a morula.

**Figure 1.**
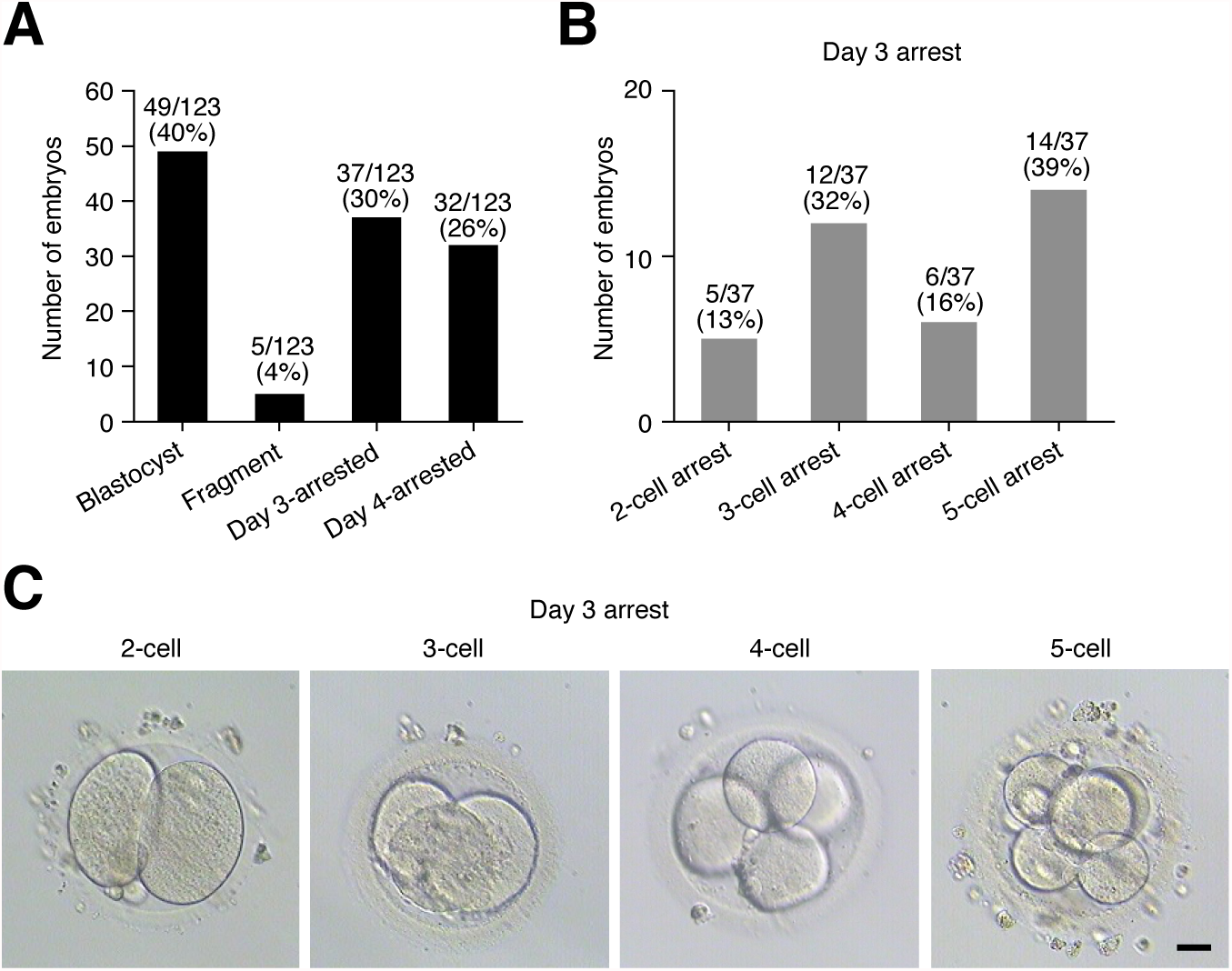
Human IVF embryos have poor *in vitro* developmental capability. **(A)** Typical outcomes for human IVF blastocyst development. Developmental results of 123 embryos were collected from 30 patients undergoing IVF. **(B)** Number of blastomeres in the 18 embryos arrested on day 3 post-fertilization. These embryos were defined as arrested and had normal morphology with distinct blastomeres, no unequal divisions and no fragmentation. **(C)** Bright-field images of uncompacted arrested human embryos between the 2-cell and 5-cell stages. Scale bar = 20 *µ*m.

To explore the mechanism of arrest, we performed single-embryo RNA-seq on 17 arrested human embryos and combined this data with 6 arrested embryos from a previous study (Sha et al., 2020). We compared these data with publicly available single cell RNA-seq or single-embryo data from normal human embryos from the oocyte through to the late blastocyst (Blakeley et al., 2015; Petropoulos et al., 2016; Yan et al., 2013). Considering the high rate of developmental arrest of human embryos, it should be noted that the “normal” dataset is likely to contain embryos that are arrested or will go on to arrest. The data set was analyzed using a hybrid single cell-RNA-seq and bulk RNA-seq pipeline based on scTE and EDASeq G/C normalization (He et al., 2021; Hutchins et al., 2017; Risso et al., 2011). In total, our dataset contained 1,020 single cells or single embryos, of which 23 were arrested.

### Arrested embryos can be divided into three types of arrest

We next investigated the developmental state of the arrested embryos. Potentially, arrested embryos have a failed or distorted developmental program, which may explain their arrest. Projection of the gene expression into PCA placed the arrested embryos in several locations, ranging from a 2-cell, 4-cell state, to 8-cell through morula (**Fig. 2A**). Overall, the arrested embryos did not diverge from a normal developmental pathway and clustered with normal embryo data from the zygote through to the late morula (E4) stage. This suggests that development is not drastically disturbed in the arrested embryos. Interestingly, all of the arrested embryos had not undergone compaction, yet many arrested embryos had a gene expression signature in advance of their morphological state. This suggests the developmental program is unlinked from compaction and cell division, and blastomeres can adopt a gene expression program unlinked from their morphological state.

**Figure 2.**
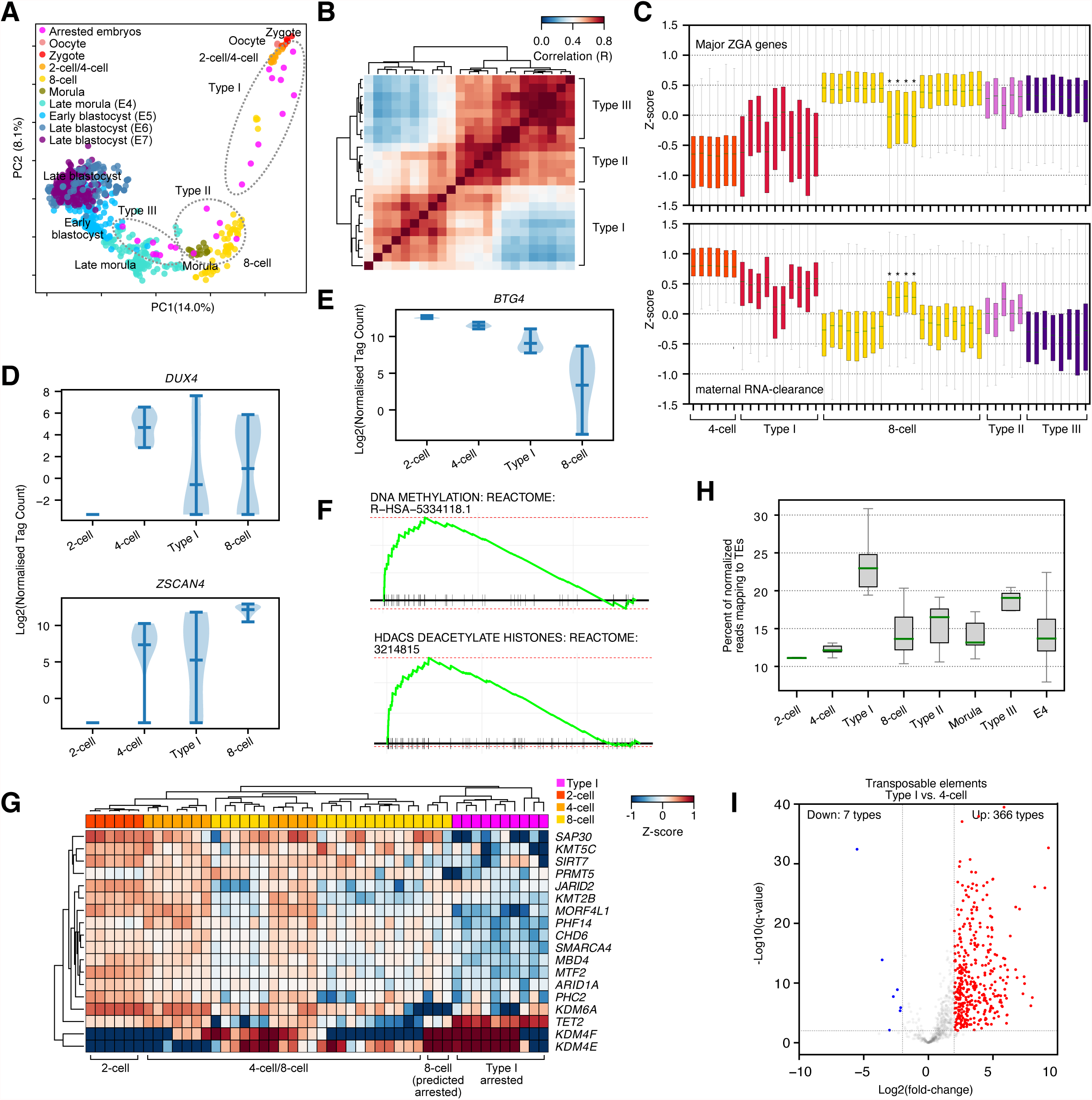
Arrested embryos adopt three distinct types of arrest. **(A)** Principal component analysis (PCA) of single embryo or single cell RNA-seq for normal or arrested human embryos. Arrested embryos are marked in pink. Presumed normal embryonic cells are colored by their developmental stage. The control (non-arrested presumed normal) samples are from a reanalysis of GSE66507 (Blakeley et al., 2015), PRJEB11202 (Petropoulos et al., 2016) and GSE36552 (Yan et al., 2013) embryonic data. E = Embryonic-stage samples, as defined in (Petropoulos et al., 2016), for this and all subsequent figures. **(B)** Pair-wise co-correlation matrix (Pearson’s R) of the arrested embryo RNA-seq data. The types (based on the major clades in each cluster) are indicated. Clustering is based on Euclidean distance with complete linkage and optimal ordering. **(C)** Box plots showing the expression in each cell/embryo for the major ZGA genes and maternal RNA clearance genes as defined in **Supplementary Fig. 3A**). **(D)** Violin plot showing the expression of the key ZGA genes *DUX4* and *ZSCAN4*. **(E)** Violin plot of expression for the key maternal RNA clearance gene *BTG4*. **(F)** Gene Set Enrichment Analysis (GSEA) showing significantly enriched gene set terms for the up-regulated genes in the Type I arrested embryos versus 8-cell stage embryos. **(G)** Heatmap of the Z-scores of the expression of a selection of differentially expressed epigenetic factors. Cells/embryos were clustered according to the genes or cells by Euclidean distance and complete linkage. Cell types are labelled in the colored header bar and labelled below the heatmap. Note that the ‘normal’ 8-cell from the (Yan et al., 2013) dataset, which appears to be failing the MZT and we predict is arrested, is indicated with stars. **(H)** Box plots showing the percent of normalized tags mapping to transposable elements in the indicated embryonic stages or the arrested embryos. **(I)** Volcano plot of differential gene expression for Type I arrested embryos versus 4-cell stage embryos. This plot only shows the differentially expressed TEs. The x-axis is the log2 fold-change, and the y-axis is the -log10(q-value) as reported by DESeq2 with Bonferroni-Hochberg multiple testing correction. Significantly up and down-regulated TEs are labelled in red and blue, respectively, and the number of TE types that are up or downregulated are labelled.

Early embryonic development is a highly dynamic process with rapid changes in gene expression (Xia and Xie, 2020; Xu et al., 2021). The arrested embryos are distributed through several developmental stages, hence to isolate the factors involved in arrest we must remove development as a confounding variable. Co-correlation of the arrested embryos resulted in three groups (**Fig. 2B**), which we designate Type I-III. Type I embryos clustered with the zygote, 2-cell and 4-cell stages, whilst Type II and III were grouped with 8-cell and beyond stages, (**Fig. 2A**). Analysis of the arrested embryo types using CytoTRACE, which estimates developmental trajectories (Gulati et al., 2020), placed Type I between 2-cell and 4-cell, Type II between 4-cell and 8-cell, and Type III between E3 (Embryonic-stage 3, early morula, as defined in (Petropoulos et al., 2016)) and morula stages (**Supplementary Fig. 1A-C**). Interestingly, CytoTRACE did not indicate that the arrested embryos had lost developmental potency, supporting the idea that the developmental gene expression program is not substantially impacted by arrest. Analysis of genes specific to each stage of development indicated that each arrested Type expressed genes normal for their developmental stage (**Supplementary Fig. 1D-F**). For example, Type I arrested embryos expressed typical developmental markers for the 4/8-cell stages, such as *LIN28A, DIS3, REST* and *SNAPC1* (**Supplementary Fig. 1D**). Whilst Type II and III arrested embryos expressed markers specific for the morula or even blastocyst, including *NANOG, DNMT3L, ESRRB*, and *ZFP42* (**Supplementary Fig. 1E**). Clustering the expression of genes from an 8-cell-specific signature identified in (Hasegawa et al., 2015) correctly clustered Type II and III with 8-cells, and Type I with 2/4-cell stages (**Supplementary Fig. 1F**). This suggests that the Type I arrested embryos are developmentally arrested at the 2/4-cell stage, whilst Type II and III embryos are more developmentally advanced and express 8-cell, morula and even blastocyst-stage developmental genes.

### Arrested embryos do not show excessive aneuploidy

We next looked at the karyotype of the arrested embryos. Normal human embryos have surprisingly high levels of aneuploidy, compared to other species (Vanneste et al., 2009), and this may be a contributing factor to arrest during IVF, particularly in embryos from women of advanced maternal age (Qi et al., 2014). Aneuploidy mainly takes two forms (Zamani Esteki et al., 2019): full embryo aneuploidy due to meiotic errors in the oocyte/zygote, or mosaic aneuploidy due to mitotic errors after fertilization (McCoy, 2017; Vera-Rodriguez et al., 2015). We used the RNA-seq data to estimate the karyotype, as we could then correlate it with the arrest type in the same embryo. We used the method described in (Griffiths et al., 2017) to estimate aneuploidy. Around 30% of cells were predicted to be aneuploid (**Supplementary Fig. 2A, B**), which agrees with previous data that aneuploidy is common in human embryos (Thomas et al., 2021). We notice that chromosomal defects become particularly evident at the 8-cell stage, and reach around 30% at the morula/E3 stage (14/48 cells, 29%) and persist to the E7-stage (late blastocyst) at similar rates (116/321 cells, 36%) (**Supplementary Fig. 2C**). Of the arrested embryos, 6/23 (26%) had a predicted aneuploidy (**Supplementary Fig. 2A, B**). This number is not substantially different from normal embryos and is in line with the typical levels of aneuploidy seen in human embryos. This computational approach cannot easily detect the difference between meiotic and mitotic aneuploidies, but assuming most of the aneuploidies we see are due to mitotic errors, there was no overall bias in the gain or loss of specific chromosomes (**Supplementary Fig. 2C, D**). Ultimately, these data suggest that aneuploidy is not a specific feature of arrested embryos. In a study of aneuploidy in embryos from women of advanced maternal age, 50% of embryos still developed to the blastocyst stage, despite 84% of the embryos having at least one chromosomal abnormality (Qi et al., 2014). Similarly, there is evidence that mosaic aneuploidies are common and may not be detrimental to development (McCoy, 2017; Starostik et al., 2020), at least to the blastocyst stage. Finally, meiotic aneuploidies can develop to the blastocyst stage, although they have severe consequences for further development (Shahbazi et al., 2020). Hence, we argue that whilst aneuploidy is an important problem in post-implantation development, it is not a dominant factor for developmental competency to reach the blastocyst, and it is not responsible for developmental arrest pre-compaction.

### Type I arrested embryos fail the maternal-to-zygotic transition

We next looked at specific biological processes that were behind the arrest. Because each type of arrested embryo clusters with different cell types, we decided to investigate each type separately. The arrest of Type I embryos is developmentally close to the maternal-to-zygotic transition (MZT), which happens from fertilization to the 8-cell stage (Vastenhouw et al., 2019). The MZT encompasses two processes, major zygotic genome activation (ZGA), and the degradation of maternal transcripts. A failure of the major ZGA or maternal-clearance may lead to embryonic arrest. Indeed, there is evidence that maternal-clearance is defective in arrested embryos (Sha et al., 2020), but the major ZGA can initiate normally in arrested embryos (Dobson et al., 2004). We first defined major ZGA genes as those genes that were significantly up-regulated >4-fold from 2-cell to 8-cell stages, and maternal-clearance genes as those significantly down-regulated (**Supplementary Fig. 3A and Supplementary Table 1**), in a method similar to (Gao et al., 2018). Gene set enrichment analysis (GSEA) supported the designation of these genes as representing the MZT, as up-regulated terms included spliceosomes, transcription, and down-regulated genes included female gamete generation (**Supplementary Fig. 3B**).

We next applied these two MZT gene sets to the arrested embryos. Surprisingly, the expression of major ZGA genes and maternal-clearance genes could discriminate Type I arrested embryos from Type II and III (**Fig. 2C**). Type II and III arrested embryos had gene expression levels of major ZGA genes that matched the 8-cell stage, and maternal-clearance genes were lower in type II/III than in 4-cell-stage embryos (**Fig. 2C**), indicating that Type II and III arrested embryos had traversed the MZT. Conversely, the Type I arrested embryos had poor activation of ZGA genes and incomplete degradation/reduction of maternal transcripts (**Fig. 2C**). This observation was supported by looking at the expression of key MZT-related regulatory genes. The DUX family of transcription factors are involved in ZGA, and *DUX4* is activated just before major ZGA (De Iaco et al., 2017; Hendrickson et al., 2017). In the Type I arrested embryos *DUX4* remained low, compared to the expression of *DUX4* in 4-cell stage embryos, and two target genes of DUX4, *DUXA* and *ZSCAN4* were poorly induced (**Fig. 2D, and Supplementary Fig. 3C**). Similarly, the key maternal RNA-clearance genes *BTG4, PAN2* and *CNOT6L* (Yu et al., 2016), remained high in Type I arrested embryos (**Fig. 2E, and Supplementary Fig. 3D**). Interestingly, our data indicates that one of the ‘normal’ 8-cell embryos resembles a type I arrested embryo, with low levels of major ZGA genes and incomplete maternal RNA-degradation (**Fig. 2C**). Overall, our data suggest that ZGA failure can account for about ∼40% of arrested embryos (10/23 arrested embryos and 1/4 ‘normal’ embryos). Overall, these results suggest that a failure to traverse the MZT is responsible for the arrest of Type I embryos.

We next looked at mechanisms underlying the arrest of Type I embryos. The MZT is a time of very active epigenetic and 3D genome rearrangements (Xia and Xie, 2020), and we speculated that epigenetic defects underlie MZT. We first measured differentially regulated genes and transposable elements (TEs) by comparing Type I versus 4-cell stage embryos (**Supplementary Fig. 4A**). GSEA indicated that DNA methylation and HDAC deacetylase pathways were upregulated (**Fig. 2F**), suggesting epigenetic regulatory dysfunction. Many specific epigenetic repressors and activators were significantly downregulated in arrested embryos (**Fig. 2G**). For example, *SAP30*, a member of the SIN3A co-repressor complex was downregulated, along with the histone arginine methyltransferase *PRMT5*. Histone lysine methylating enzymes were also downregulated, including *KMT5C (SUV420H2*), which is responsible for catalyzing the repressive histone H4K20me3 mark. Additionally, *KMT2B (MLL2*), an enzyme responsible for the active histone mark H3K4me3 was also downregulated. Conversely, the histone demethylases *KDM4F* and *KDM4E* were up-regulated in arrested embryos. The consequences of these changes are likely to be major disruptions in the balance of activatory and repressive chromatin.

Ideally, we would perform ChIP-seq for methylated histones in human embryos. However, this technique is extremely challenging when small amounts of material are available. Histone ChIP-seq has been performed using mouse embryos but required several hundred embryos (Liu et al., 2016), which is a level of material not available for human embryos. Consequently, we exploited an indirect method to measure the epigenetic state of the cell. In other embryonic model systems, such as mouse or human embryonic stem cells (ESCs), we have and others have shown that when epigenetic regulators are disrupted, transposable elements (TEs) tend to be upregulated (He et al., 2019; Matsui et al., 2010). The situation is somewhat complicated by the normal presence of TEs during embryonic development as TE sequence fragments are expressed during early embryonic development (Goke et al., 2015). Nonetheless, TE fragment expression can act as an indirect read-out for chromatin state. Remarkably, in the Type I arrested embryos, we observed a dramatic increase in the overall number of reads mapping to TEs (**Fig. 2H**). This was also reflected in the number of differentially regulated TEs, and 366 TE types were up-regulated, whilst only 7 types of TE were downregulated (**Fig. 2I**). This pattern held whether we compared Type I to 2-cell, 4-cell or 8-cell cells (**Supplementary Fig. 4B**), and TEs were not appreciably differentially regulated in Type II and Type III arrested embryos, which also had low levels of TE mapped reads (**Fig. 2H and Supplementary Fig. 4C**). Of the types of deregulated TEs, endogenous retrovirus (ERV) families were up-regulated, and ∼70% of ERV1, ERVL, and ERVK family TEs were upregulated (**Supplementary Fig. 4D**). Only a few SINE families were upregulated, but LINE:L1s were upregulated, including the L1HS family of TEs that are capable of transposition (Beck et al., 2010) (**Supplementary Fig. 4E**). Indeed, a major epigenetic suppressor of LINE L1s, PRMT5 (Kim et al., 2014), was significantly down-regulated in Type I arrested embryos (**Fig. 2G**). Ultimately, the massive deregulation of TEs seen in arrested embryos is reminiscent of the widespread activation of TE families in response to the knockdown of epigenetic regulators we previously observed in mouse ESCs (He et al., 2019). One further point to note is that many of the deregulated epigenetic factors in Type I embryos are repressors (**Fig. 2G**). This may seem counterintuitive that repressors are downregulated, TEs are upregulated, but the ZGA fails to activate. However, epigenetic repression is an important, if incompletely understood, component of the ZGA (Wu et al., 2018). Overall, our data suggest that Type I arrested embryos have MZT defects that are likely due to epigenetic deregulation.

### Type II and III arrested embryos enter a quiescent-like state

We next looked at Type II and III arrested embryos. These arrested embryos are distinct from Type I, by overall gene expression patterns (**Fig. 2B**), successful traversal of the MZT (**Fig. 2C**), and do not appear to have a deregulated epigenetic state, as measured by TE sequence fragment expression (although some LINE L1s are upregulated) (**Supplementary Fig. 4C, E**). This suggests other mechanisms are responsible for Type II and III arrest. Embryonic development is a highly dynamic process, hence we needed to determine the most appropriate developmental stage to compare arrested embryos. PCA indicates the Type II and III arrested embryos are most similar to cells of the 8-cell/morula stage and the late morula (E4)-stages (**Fig. 2A**). To determine the closest comparable embryonic stage, we measured the number of significantly differentially expressed (DE) genes by comparing Type II and III embryos to 8-cell, morula, and E4-stages (**Supplementary Fig. 5A**), and chose the comparison with the smallest overall number of differentially regulated genes. This approach suggested that the closest comparisons are Type II versus morula (1034 DE genes), and Type III versus E4 (876 DE genes) (**Supplementary Fig. 5A, B and Supplementary Table 2**).

We next analyzed the sets of genes that were deregulated for Type II and III. Interestingly, for both types, ribosomes, histones and translation-related genes were downregulated, as determined by GSEA and gene expression levels (**Fig. 3A-C**). Ranking cells by the sum of expression of small and large ribosomes and nucleosomes placed almost all arrested embryos at the bottom of the list (**Supplementary Fig. 5C-E**), as exemplified by the expression of selected histone and ribosome genes (**Supplementary Fig. 5F, G**). Based on the downregulation of nucleosomes and ribosomes, we reasoned that the arrested embryos were entering into a quiescent or senescent-like state. This was supported by GSEA, which suggested a senescence/quiescence gene expression program was being activated (**Fig. 3D**). We used the RNA-seq data to estimate the cell cycle phase/activity, based on specific marker genes (Macosko et al., 2015), and this analysis suggested that all except two of the embryos had low levels of cell cycle activity (**Fig. 3E and Supplementary Fig. 6A, B**). Expression of the A-type cyclin *CCNA2* was downregulated in arrested embryos, whilst the cell cycle inhibitor CDNK1A (p21) was upregulated (**Fig. 3F**). Similarly, the negative cell cycle regulator *RB1* was expressed at higher levels in arrested embryos (**Fig. 3G**). Other genes positively involved in the cell cycle were similarly downregulated in arrested embryos, such as the tubulin subunits *TUBB4B, TUBA1A, CENPA and BCLAF1*, a negative regulator of p21 expression (Lee et al., 2012) (**Supplementary Fig. 6B, C**). These results suggest that the Type II and III arrested embryos are entering into a quiescent/senescent-like state marked by reductions in ribosomes, nucleosomes, protein translation and cell cycle factors.

**Figure 3.**
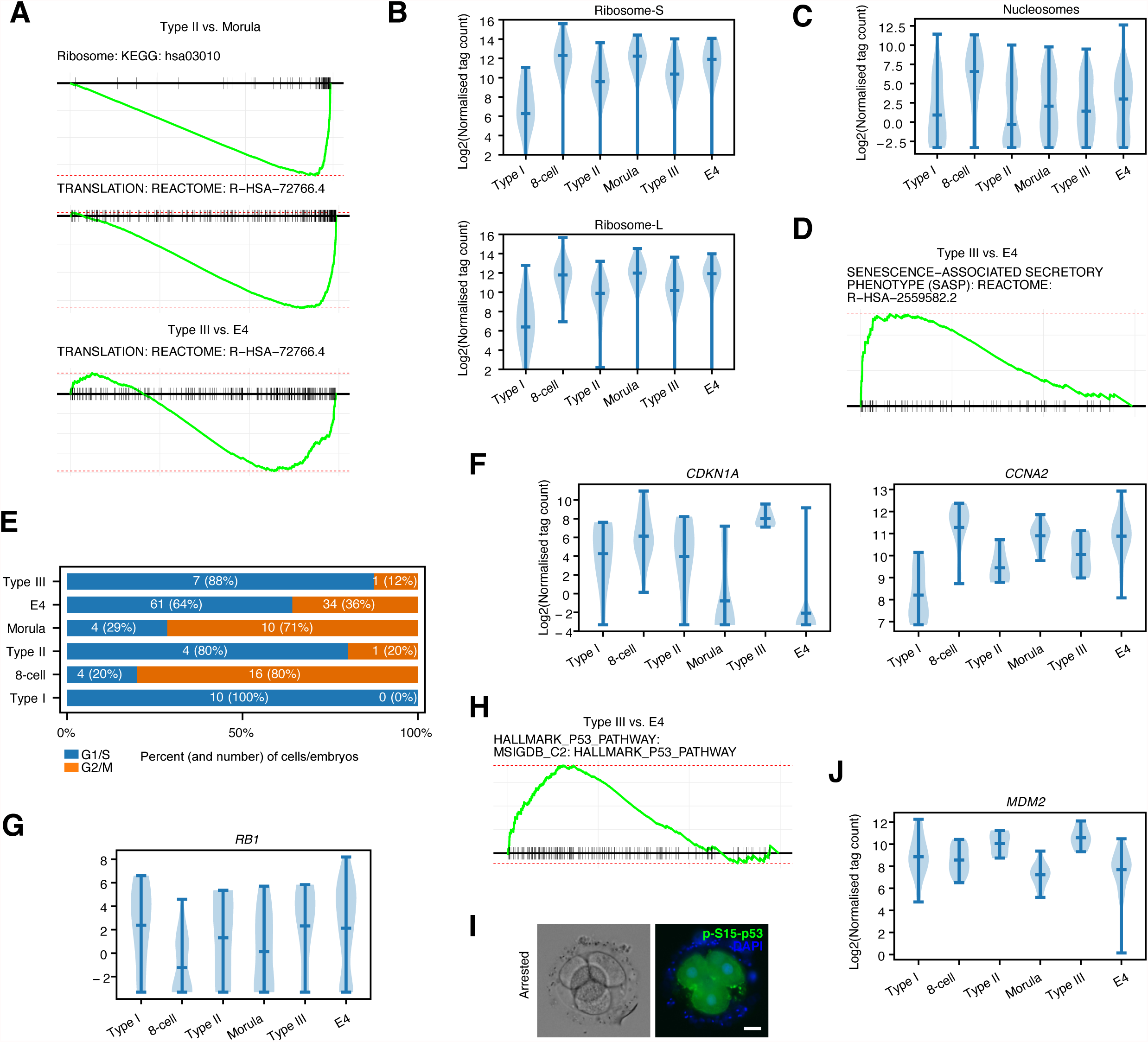
Arrested embryos downregulate histones, ribosomes and adopt a quiescent/senescent-like gene expression program. **(A)** GSEA for down-regulated genes in Type II versus morula comparison. **(B)** Violin plot of expression for all small (top) and large (bottom) ribosomal subunits in the indicated embryonic stages or arrested embryos. **(C)** Violin plot of expression for all expressed nucleosomes in the indicated embryonic stages or arrested embryos. **(D)** GSEA showing the SASP (senescence-associated secretory phenotype) term is significantly enriched in Type III versus E4 (late morula) comparison. **(E)** Cell cycle analysis of the 8-cell normal or arrested embryos. Selected G1/S or G2/M genes are shown in **Supplementary Fig. 6A**. This panel should be interpreted as more of a measure of the cell cycle activity for each class of embryo, rather than a definitive call for each cell in the indicated cell cycle phase. **(F)** Violin plot showing the expression of *CDKN1A* (p21), or *CCNA2* in the indicated embryonic stages or arrested embryos. **(G)** Violin plot showing the expression of *RB1* (Retinoblastoma1) in the indicated embryonic stages or arrested embryos. **(H)** GSEA showing the enrichment of the HALLMARK p53 list of target genes is significantly enriched in Type III versus E4 (late morula) comparison. **(I)** Phospho-S15-p53 (green) immunostaining in arrested embryos. Embryos are co-stained with DAPI (blue), scale bar = 20 *µ*m. **(J)** Violin plot of expression showing the expression level of the p53 target gene *MDM2* in the indicated embryonic stages or arrested embryos.

Quiescence is a defined molecular program (Coller, 2011), but the details remain elusive and the mechanism is not fully elaborated as quiescence as a state is diverse and exists in many cell types within the somatic body and during development (Marescal and Cheeseman, 2020). One key regulator identified in hematopoietic stem cells (HSCs) is p53 (Liu et al., 2009). GSEA of up and down-regulated genes suggested hyperactivation of p53 target genes (**Fig. 3H**), which is also a mark of HSC quiescent cells (Liu et al., 2009). The mRNA level of p53 (*TRP53*) was relatively unchanged (**Supplementary Fig. 6D**). However, immunofluorescence indicated that arrested embryos had high levels of phosphorylated (Ser15) p53 (**Fig. 3I**). The major target gene of p53, *MDM2*, was also upregulated (**Fig. 3J**), a feature also seen in quiescent HSCs (Liu et al., 2009). Finally, ranking all cells by the sum of their gene expression for p53 target genes (i.e. doing the inverse of a GSEA) placed all arrested embryos at the top of the list (**Supplementary Fig. 6E**), indicating increased activity downstream of p53. This analysis suggests that the arrested embryos are becoming quiescent, with a deregulated cell cycle and activated p53. However, the developmental program is still being executed, which suggests that development and cell cycle are relatively uncoupled processes.

### Partial rescue of arrested embryos by resveratrol

The arrested embryos have an arrested cell cycle. However, it is unclear if the arrest is related to quiescence (i.e. reversible), or senescence (i.e. irreversible). Senescence and quiescence have many similar cellular and mechanistic features; and there is no clear way to discriminate between the two states, except for reversible reactivation (Terzi et al., 2016). To determine if the arrested embryos are irreversibly arrested, we attempted to reactivate the embryos and recommence development. We selected several small molecule inhibitors that have previously been shown to impact embryonic or pluripotent stem cell development. Specifically, the mTOR inhibitor rapamycin, ERK inhibitor PD0325901, Vitamin C, and resveratrol. These four compounds have been implicated in various aspects of senescence/quiescence and embryogenesis. Rapamycin inhibits mTOR to promote autophagy and has been previously shown to improve pig oocyte development (Elahi et al., 2017). Vitamin C is an antioxidant and epigenetic modulator that can affect DNA methylation by functioning as a co-factor for DNA demethylation TET enzymes (Blaschke et al., 2013). ERK inhibition assists in the establishment of the inner cell mass in the blastocyst stage (Van der Jeught et al., 2013). Finally, resveratrol is an anti-oxidant that can improve pig and bovine oocytes (Itami et al., 2015; Pasquariello et al., 2020; Takeo et al., 2014), and *in vitro* development of aged mice and human oocytes (Liu et al., 2018).

Application of rapamycin, PD0325901, and vitamin C to the arrested embryos had only a limited impact, and the embryos did not recommence development at rates substantially higher than control (untreated arrested) embryos (**Supplementary Fig. 7A**). Only resveratrol, an anti-oxidant with SIRT activating effect had a substantial impact on the development of arrested embryos. After treatment, 23/42 embryos recommencing development (**Fig. 4A**). However, it should be noted that 4/42 embryos reinitiated development, but ultimately fragmented, suggesting that at least some of the reactivated embryos are incapable of, or unsuitable for, further development. Similarly, whilst many (19/42, 45%) of the embryos recommenced development, only 9 compacted, and only 3 made it to the blastocyst stage (**Fig. 4B, C**). A caveat should, however, be applied. The arrested embryos were cultured for a further day before starting treatment with resveratrol (i.e. the day 3-arrested embryos started treatment on day 4). Potentially, treating the embryos with resveratrol at an earlier time point may have activated more embryos without disintegration.

**Figure 4.**
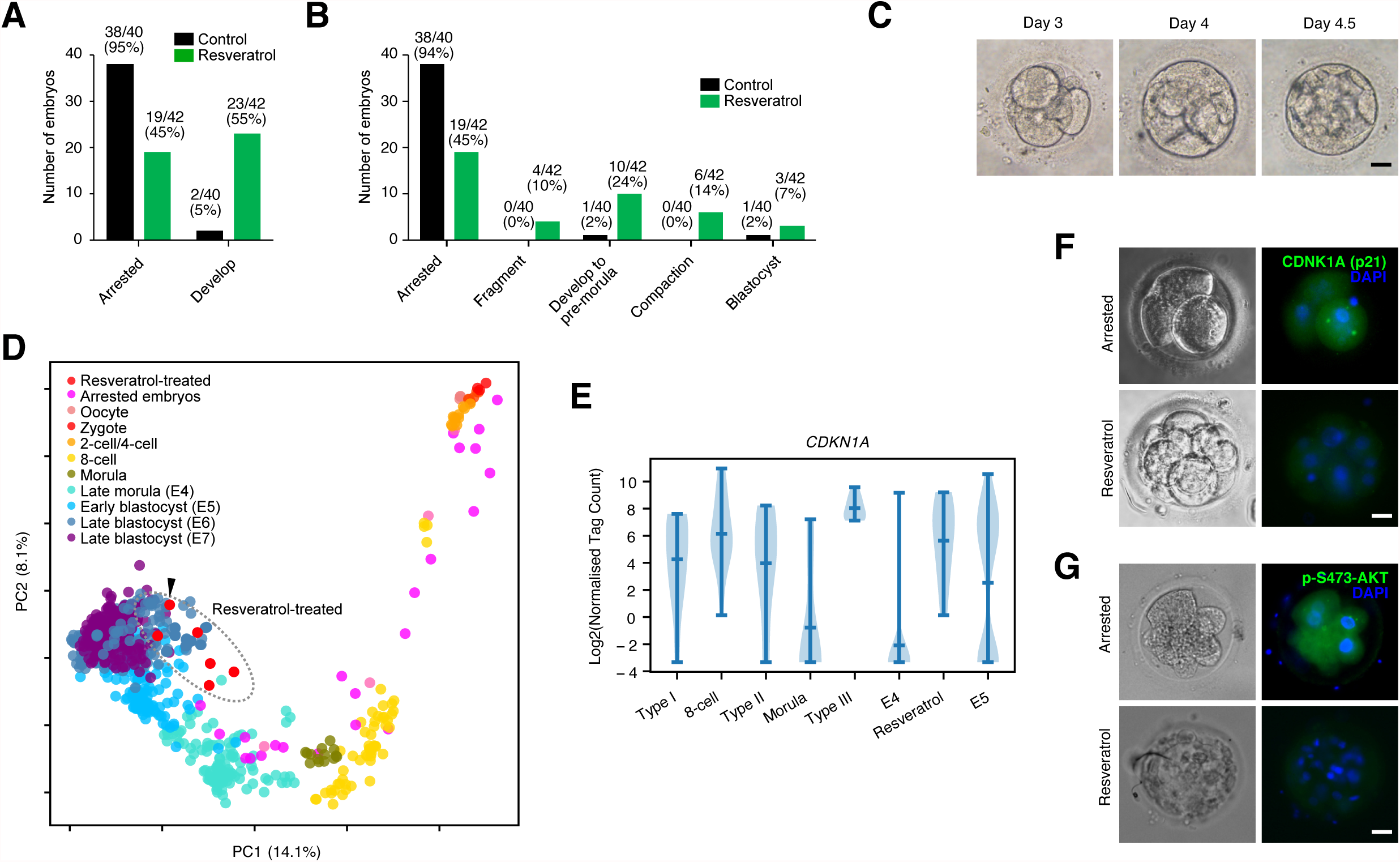
Resveratrol can overcome embryonic arrest in limited cases. **(A)** Number of embryos that remain in an arrested state, or recommenced development (including those embryos that fragment). Embryos were either left untreated or treated with resveratrol. The total number of embryos in the control group is 40, and 42 in the resveratrol group. **(B)** Bar chart showing the breakdown of the stages of embryonic development the control and resveratrol embryos reached. **(C)** Morphology of a reactivated resveratrol-treated embryo. The embryo was arrested on day 3 and did not show any degeneration at day 5, and then treated with resveratrol and proceeded to the early blastocyst-like stage. Scale bar = 20 *µ*m. **(D)** PCA of normal human embryo RNA-seq, with arrested embryos (pink) and embryos treated with resveratrol (red). The embryo in **panel C** is marked with a black arrow. **(E)** Violin plot showing the expression of *CDNK1A* in the indicated embryonic stages arrested embryos or the reactivated embryos treated with resveratrol. **(F)** Immunostaining and brightfield of embryos either untreated or treated with resveratrol. Immunofluorescence using an antibody against p21 (*CDKN1A*) (green). Embryos are co-stained with DAPI (blue), scale bar = 20 *µ*m. **(G)** As in **panel F**, but for phospho-S473-AKT (green). Embryos are co-stained with DAPI (blue), scale bar = 20 *µ*m.

To further investigate the gene expression patterns, we performed single-embryo RNA-seq on the resveratrol-treated embryos. PCA suggested that the resveratrol-treated embryos had indeed recommenced development (**Fig. 4D**), as the resveratrol-treated embryos were now grouped with early and late blastocyst-stage embryos, and normal developmental marker genes were activated. For example, *ESRRB* and *TFCP2L1* were highly expressed (**Supplementary Fig. 7B**). We next looked at the quiescence-related genes. Curiously, ribosomes and nucleosomes had not recovered to their normal levels (**Supplementary Fig. 7C**), nor had the quiescence-related genes *CENPA* and *RB1* identified in the arrested embryos (**Supplementary Fig. 7D**). Expression of the cell cycle inhibitor *CDKN1A* had also not declined (**Fig. 4E**). However, immunostaining of arrested embryos and embryos treated with resveratrol indicated that the protein levels of p21 (*CDKN1A*) were reduced, suggesting resveratrol is affecting p21 post-translationally (**Fig. 4F**). Similarly, phosphorylated AKT (Ser473) was also reduced (**Fig. 4G**), suggesting reduced cross-talk between RB and AKT, which is a hallmark for reduced quiescence/senescence in mouse liver cells (Imai et al., 2014). Overall, resveratrol partially reactivates arrested embryos, although they retain high expression of quiescent-markers, but low protein levels. Resveratrol can reactivate a minority of embryos, which activate a normal-like developmental program and a subset of embryos can make it to the blastocyst. However, the resveratrol-treated embryos still have deregulated levels of ribosomes, protein translation, and have not adopted a completely normal state.

### SIRT activators resveratrol and nicotinamide riboside affect embryonic metabolism

Resveratrol has two main mechanistic functions: As an antioxidant, and also as a co-activator of the NAD+ dependent deacetyltransferase SIRT1 (Borra et al., 2005). RNA-seq analyses showed that the expression *SIRT1* RNA was low in Type I-arrested embryos, but was high in Type II/III and resveratrol-treated arrested embryos (**Fig. 5A**). Using immunofluorescence, SIRT1 protein levels were low in the arrested embryos, but when treated with resveratrol, SIRT1 levels increased (**Fig. 5B**). We also measured NRF2 protein levels (**Fig. 5C**), as resveratrol has been reported to confer its antioxidant benefits by upregulating NRF2 at both the transcriptional and protein levels (Ungvari et al., 2010). Resveratrol indeed activated the antioxidant protein NRF2. These data suggest that resveratrol is activating both SIRT and an anti-oxidant effect, although it is not clear which of the two are important for the reactivation of the arrested embryos.

**Figure 5.**
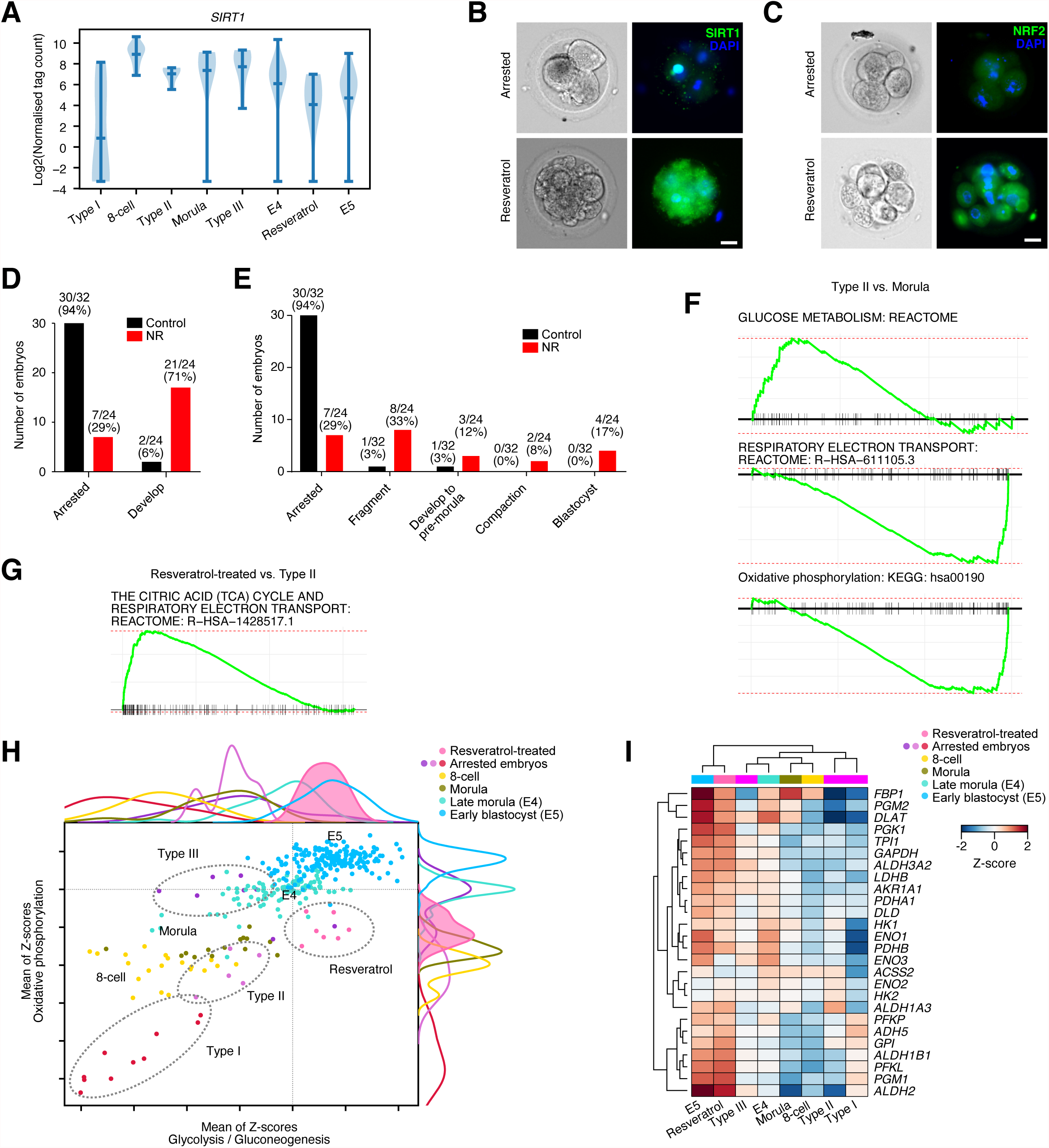
Resveratrol and nicotinamide riboside reactivates development partly through modulation of SIRT activity. **(A)** Violin plot showing the expression of SIRT1 in the indicated embryonic stages or arrested embryos or resveratrol-reactivated embryos. **(B)** Immunofluorescence staining of SIRT1 (green) and brightfield in arrested or resveratrol-reactivated embryos. Embryos are co-stained with DAPI (blue), scale bar = 20 *µ*m. **(C)** Immunofluorescence staining of NRF2 (green) and brightfield in arrested or resveratrol-reactivated embryos. Embryos are co-stained with DAPI (blue), scale bar = 20 *µ*m. **(D)** Number of embryos that remain in an arrested state, or recommenced development (including those embryos that fragment). Embryos were either left untreated or treated with NR (nicotinamide riboside). The total number of embryos in the control group is 32, and 24 in the NR-treated group. **(E)** Bar chart showing the number of arrested embryos that would reinitiate development (including embryos that would ultimately fragment). The total number of embryos in the control group is 32, and 24 in the NR-treated group. **(F)** GSEA for up and down-regulated genes in Type II versus morula comparison. **(G)** GSEA for up and down-regulated genes in resveratrol versus Type II arrested embryos. **(H)** 2D dot plot showing the sum of the Z-scores for the genes in the indicated KEGG categories. The x-axis scores the glycolysis/gluconeogenesis pathway, the y-axis the oxidative phosphorylation pathway. Each dot in the plot is a cell/embryo, and the top and right axis have the kernel density for each group of cells. The resveratrol-treated embryos have a filled-in color (pink) for emphasis. The locations of the arrested and resveratrol-treated embryos are indicated by dashed lines and labels, and the normal developmental states are indicated by labels. **(I)** Heatmap showing the expression of select glycolytic metabolic genes.

The antioxidant vitamin C failed to reactivate arrested embryos (**Supplementary Fig. 7A**), which suggests an anti-oxidant effect is not the main factor for embryo reactivation. Hence, resveratrol may be reactivating development by modulating SIRTs. We next treated arrested embryos with a second SIRT activator NR (Nicotinamide Riboside), that does not have reported anti-oxidant capability. NAD+ availability is a rate-limiting step for SIRT activity, and SIRTs can be activated by NR which is converted to NAD+ and increases the activity of SIRT1 and SIRT3 in cells (Canto et al., 2012). When arrested embryos were treated with NR, in an effect similar to resveratrol, they were reactivated and would proceed through development, and a few embryos reached a blastocyst-like state (**Fig. 5D, E**). As NR and resveratrol can phenocopy, it suggests that the antioxidant activity of resveratrol is not required, and the dominant pathway in the reactivation of arrested embryos is the activation of SIRTs.

SIRT enzymes have a key role in controlling the balance of glycolytic, oxidative and fatty acid metabolic processes in somatic cells (Canto et al., 2012; Chang and Guarente, 2014). The developing human embryo undergoes significant changes in the metabolic pathways utilized at each embryonic stage. However, these metabolic changes remain somewhat unclear due to the difficulty of directly assaying metabolic products in small numbers of cells (Zhang et al., 2018). Briefly, from the zygote to the morula, embryos use a form of oxidative metabolism based on pyruvate. In the pre-implantation blastocyst, embryos convert to a balanced glycolytic/oxidative phosphorylation-based metabolism using glucose as a fuel source, before transitioning to glycolysis in the low oxygen environment after implantation (Zhang et al., 2018). Studies in human and mouse naïve and primed PSCs, that resemble the pre-implantation and the post-implantation epiblast respectively (Sun et al., 2021), suggest that HIF1A and the balance between SIRT1 and SIRT2 activity is important (Cha et al., 2017; Zhou et al., 2012). Reduced SIRT1 activity leads to reduced glycolysis in both embryonic and somatic cells (Cha et al., 2017; Ryall et al., 2015). To explore metabolism in the arrested embryos, we would ideally measure the metabolic products directly. However, this is infeasible with current technology, which requires hundreds or thousands of embryos for mass spectrometry-based approaches (Chi et al., 2020; Li et al., 2020). Hence, we attempted to infer metabolic state based upon the RNA-seq data. This approach is fraught with difficulty as metabolic enzymes generally have high steady-state levels of mRNA that do not respond to changes in metabolism. Consequently, we infer metabolism based on the changes in RNA levels of sets of metabolic-associated transcripts. GSEA indeed suggested that glucose metabolism was upregulated and oxidative phosphorylation was downregulated in Type II arrested embryos (**Fig. 5F**). GSEA for resveratrol-treated embryos also supported increased expression of oxidative phosphorylation genes, when we compared the resveratrol treated embryos to Type II or III arrested embryos (**Fig. 5G, and Supplementary Fig. 8A, B**). We employed a novel technique that used the sum of Z-scores for a set of genes to infer pathway activity. Plotting the sum of Z-scores of oxidative phosphorylation versus glycolysis/gluconeogenesis gene sets (from Reactome) suggested that resveratrol was pushing cells towards an increased glycolytic metabolism, whereas all the arrested cells had reduced glycolysis (and variable levels of oxidative phosphorylation) (**Fig. 5H**). This is illustrated by the RNA levels of glycolytic-related genes, such as *ALDH1B1, GAPDH, PGM1* and *PFKL*, which were higher in resveratrol-treated embryos (**Fig. 5I and Supplementary Fig. 8C**). It should be noted though, that resveratrol had not returned the reactivated embryos to a completely normal expression state, and there remain several hundred genes that are significantly differentially expressed between resveratrol-treated and E5 (early blastocyst) embryo cells, including many biological pathways that remain low (**Supplementary Fig. 8D, E**).

Finally, in our analysis of metabolism, we noticed that resveratrol also upregulated fatty acid metabolism-related genes (**Supplementary Fig. 8B, S9A, B**). This is reminiscent of the situation in mouse hepatocyte cells, where the loss of *Sirt1* leads to reduced fatty acid oxidation, due to deregulation of PPAR-genes (Purushotham et al., 2009), and agrees with several studies that show resveratrol activates SIRT1 to decrease lipogenesis and increase fatty acid oxidation in many cellular contexts (Ding et al., 2017). Overall, our data suggest that arrested embryos erroneously maintain an oxidative phosphorylation-biased metabolism and fail to upregulate glycolysis. Treatment with SIRT activators can overcome this block and force the embryos to become glycolytic, exit quiescence and recommence development (**Supplementary Fig. 9C**).

### Inferred transcriptional regulation in arrested human embryos

We next took a different approach to explore transcriptional regulation in the arrested embryos. The direct assay of transcription factor (TF) binding to the genome is currently impractical in human embryos. To date, TF binding in single cells has only been mapped using a system that involves transposons to introduce novel insertions that can then be recovered from RNA-seq data (Moudgil et al., 2020). This technique requires transgene transfection and is thus impractical in arrested embryos. Chromatin accessibility has been performed in single cells (Buenrostro et al., 2015), however, for an individual cell the data remains extremely sparse and chromatin accessibility binding is determined by pooling the data from relatively large numbers of similar single cells to reconstruct chromatin accessibility. Hence, we reverted to an approach that takes advantage of the fact that TF binding is rich around the transcription start sites (TSSs) of differentially expressed genes (Hutchins et al., 2013; Hutchins et al., 2015). In total, we found many TF motifs enriched in the differentially regulated genes in each type of arrest (**Supplementary Fig. 10A**). To understand the pattern of transcriptional regulation we focused on TF-families that regulate quiescence/senescence and metabolism. The TF FOXO1 is a major regulator of senescence in somatic cells and works partly by downregulating MYC activity (Wilhelm et al., 2016). *FOXO1* mRNA was not significantly up or down-regulated in the arrested-embryos (**Supplementary Fig. 10B**). However, the FOXO1 TF-binding motif was significantly enriched in the promoters of all three types of arrested embryos (**Fig. 6A**). This suggests that whilst the RNA levels of *FOXO1* are unchanged, FOX-family activity in the arrested embryos helps induce quiescence. The MYC motif was enriched in Type I and III differentially expressed genes and MYC target genes were downregulated in Type II and III-arrested embryos (**Fig. 6B**). This agrees with the observation that MYC activity is low in diapause and quiescent mouse embryos (Scognamiglio et al., 2016). High levels of RUNX-family TF activity has been linked with quiescence in stem cells (Wang et al., 2010), and a RUNX-family motif was enriched in all differentially expressed genes (**Fig. 6A**). Overall, this TF binding inference suggests that MYC, FOX and RUNX families are active in the arrested embryos, and they are all markers for quiescence.

**Figure 6.**
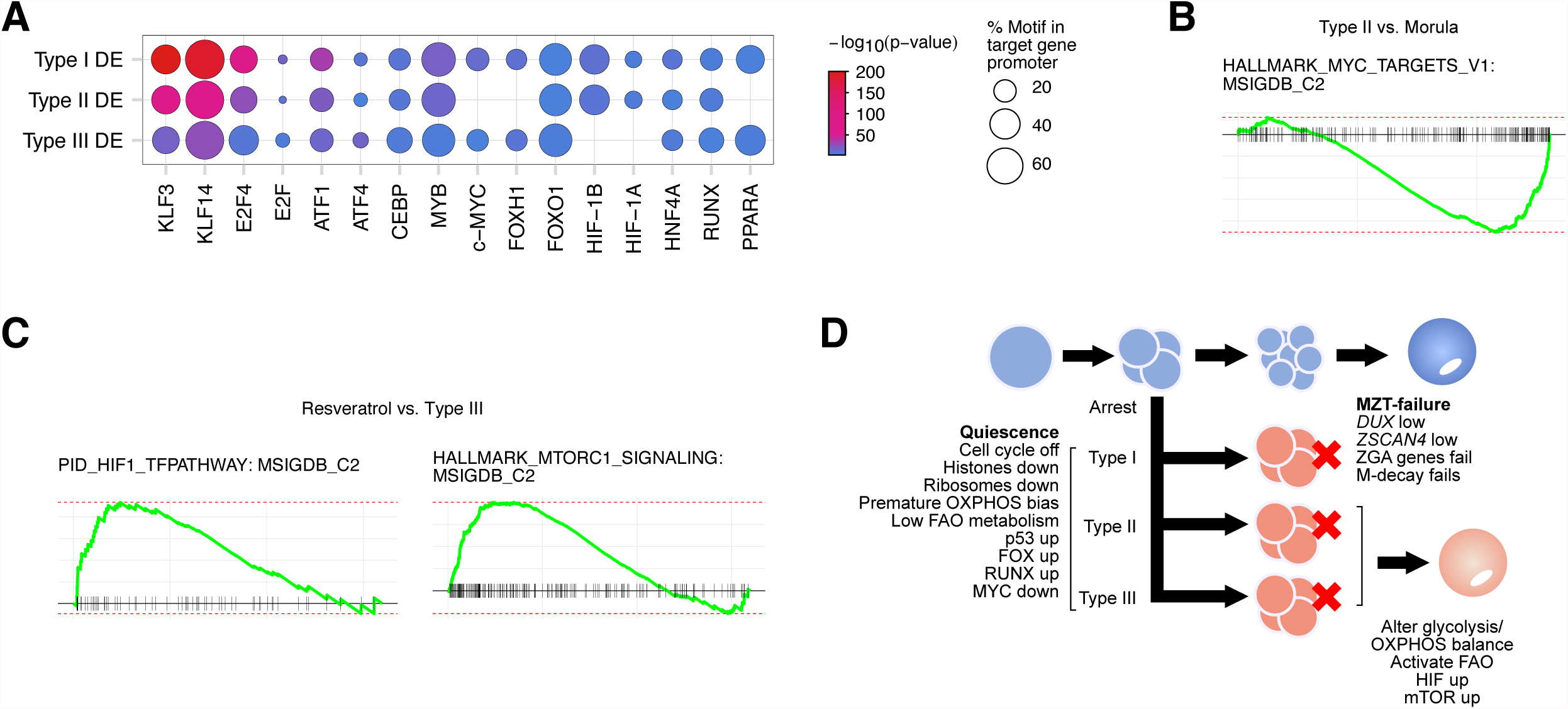
Transcriptional control of embryonic arrest. **(A)** Dot plot for enriched transcription factor motifs in the Differentially Expressed (DE) genes in the indicated arrested embryo types. Motif discovery was performed using HOMER with default settings (Heinz et al., 2010), against the promoters (defined here as −1000 bp upstream of the transcription start site) of the DE genes in the indicated types of arrested-embryo. The size of the circle indicates the percent of promoters that had the motif, and the color indicates the enrichment p-value. **(B)** GSEA of Type II vs. Morula, this panel shows the HALLMARK MYC-targets gene set. **(C)** GSEA of resveratrol-treated embryos versus Type III-arrested embryos. **(D)** Schematic model of the types of arrest and the underlying pathways driving the arrest. Type I, II and III cells all enter a quiescent state, driven by activated p53, and low MYC activity. Histones and ribosomes are downregulated and cell cycle activity is reduced. Type I arrested embryos harbor an MZT developmental error. Type II and III embryos can be induced to recommence development by modulating SIRT activity. This activates a glycolytic response and upregulation of fatty acid oxidation. OXPHOS=oxidate phosphorylation; M-decay=maternal RNA decay; FAO=fatty acid oxidation; ZGA=zygotic genome activation.

Finally, we looked at the transcriptional and signalling pathways in the resveratrol-treated arrested embryos versus the Type III arrested embryos (as the Type III arrested embryos are developmentally closest to resveratrol-treated embryos). Motif enrichment and GSEA in arrested-embryos suggested the upregulation of HIF-family TFs (**Fig. 6A**), and the HIF pathway genes were upregulated in resveratrol-treated embryos (**Fig. 6C**). This agrees with a HIF1a-driven switch from oxidative phosphorylation-based metabolism to glycolytic, as previously seen in naïve and primed mouse and human PSCs (Zhou et al., 2012). This was supported by the enrichment of PPAR-family TF motifs in the arrested differentially regulated genes (**Fig. 6A**), which is in agreement with PPAR-deregulation in *Sirt1* knockouts in mouse (Purushotham et al., 2009). Overall, the transcriptional regulation of arrested embryos suggests an upregulation of MYC activity, FOX-family TFs, possibly FOXO1, which induce quiescence.

## Discussion

Human (and primate) embryos are surprisingly poor at developing *in vitro* when compared with mouse and other mammalian species. The poor developmental capacity of human embryos has major implications for IVF. Here, we show that the arrested embryos enter a quiescent-like state. The arrested embryos downregulate histones, ribosomes, and RNA processing machinery, and upregulated p53, FOX, RUNX, and downregulated MYC activity. We classified the arrested embryos into three types based on gene expression (**Fig. 6D**). Type I appear to have failed the MZT, whilst Type II and III appear to have metabolic problems.

It should be noted that there are similarities and differences between the three types, they all arrest and acquire a quiescent-like molecular program. However, only Type I fail to correctly regulate MZT genes. The difference between Type II and III is less clear, although Type III has increased levels of genes related to oxidative phosphorylation, compared to Type I and II. Resveratrol and NR could partially overcome the arrest of the embryos, however, when we treated the arrested embryos, we did not know the type of arrest. This may help explain why resveratrol or NR can rescue only a subset of arrested embryos. We speculate that resveratrol or NR are primarily activating the Type II/III arrested embryos (**Fig. 6D**), by altering SIRT activity and metabolism. Conversely, we believe resveratrol or NR has little effect on Type I embryos, as they fail to complete the MZT, and so lack expression of key genes required for further development.

Quiescence and senescence describe the ability of some cells to exit the cell cycle. This phenomenon is best explained in somatic cells, and the mechanism is relatively well established. Stress signals, such as UV irradiation, or cellular oxidative stress lead to the deregulation of cell cycle regulatory pathways. Sometimes these activations result in senescence, which is irreversible. The state we described here for the arrested human embryos is closer to quiescence than senescence, as it is at least partially reversible. Whilst active cycling and quiescence are normal cellular responses, senescence is associated with ageing and with disease (McHugh and Gil, 2018; Terzi et al., 2016). There are, however, differences between the arrested state in the embryos and quiescence in somatic cells. We observed increased p21 (*CDKN1A*) and phospho-S15-p53, but *FOXO1* expression was not upregulated, although FOX-family TFs appear to be more active in the arrested embryos than the normal embryos. Nonetheless, there is only a partial overlap in the quiescent gene expression signature of arrested embryos and quiescent somatic cells. The downregulation of MYC activity is reminiscent of the induction of diapause in mouse blastocysts (Scognamiglio et al., 2016). Indeed, there are parallels between the MYC-induced diapause-like state, and the arrested embryos: ribosome expression is reduced, cell cycle activity declines and the cells adopt aspects of quiescence. However, diapause occurs at the blastocyst stage, whilst the arrest seen here is pre-compaction, suggesting that MYC may be involved, but the biological process is different.

Aneuploidies have been suggested to contribute to embryonic arrest. Human embryos have high levels of aneuploidy, and as much as 73% of blastocysts are mosaic (Cavazza et al., 2021; van Echten-Arends et al., 2011; Vanneste et al., 2009). However, whilst aneuploidy is highly deleterious for later post-implantation development (Hassold and Hunt, 2001), there is mixed evidence that it blocks development up to the blastocyst stage (Shahbazi et al., 2020). Indeed, meiotic aneuploid human embryos still reached the blastocyst stage (Bielanska et al., 2002), and whole chromosome aneuploidies are mainly lethal post-implantation (Shahbazi et al., 2020). Although, a screen for aneuploidy in human morula/blastocysts suggests human embryos with very high levels of mosaicism are rare (Starostik et al., 2020), suggesting a mechanism that culls severe mosaic aneuploidies before compaction (McCoy, 2017). Nonetheless, whilst we cannot rule it out, our data suggest that aneuploidy is not a dominant mechanism behind the embryonic arrest of pre-morula embryos after cleavage but before compaction.

We show that some arrested embryos can be induced to recommence development through the application of two small molecules, resveratrol and NR, both can activate the sirtuin class of deacetyltransferases. We speculate that these two drugs are altering the metabolic balance of the embryos, and are acting to push the embryos out of their quiescent-like state. Metabolism in human embryos is initially hypoxic and relies upon oxidative phosphorylation using pyruvate. Indeed, human IVF grown embryos are sensitive to high levels of oxygen (Catt and Henman, 2000), and blastulation rates are increased when oxygen was kept at 5% during embryo culture (Kovacic and Vlaisavljevic, 2008) (similar conditions were used in this study). After implantation, glycolysis becomes the dominant metabolic pathway. Our data suggest that the arrested embryos are failing to upregulate glycolytic-based or fatty-acid oxidation metabolism in advance of implantation. Overall, our data indicate two primary mechanisms to explain arrested embryos (**Fig. 6D**): A failure to correctly traverse the MZT (about 40% of arrested embryos), and a failure to rearrange metabolism towards glycolysis/fatty acid oxidation (about 60% of arrested embryos).

## Materials and Methods

### Human embryo collection and ethical approvals

All patients received standard antagonist protocol for ovarian stimulation (Yuan et al., 2021). Patients were injected with recombinant FSH (Gonal-F, Merck, Italy) on day 2 of their menstrual cycle with a starting dose of 150 IU/d. Transvaginal ultrasound and blood E2 levels were used to monitor follicle growth. When at least two follicles grew larger than 18 mm in diameter, 0.1 mg of gonadotropin releasing hormone agonist (Triptorelin, Ferring GmbH, Germany) and 4000 IU of human chorionic gonadotropin was injected as a trigger. 36 h after trigger, transvaginal ultrasound-guided oocyte retrieval was performed under anesthesia.

### *In vitro* fertilization of oocytes and culture of embryos

Fertilization and *in vitro* culture procedures were performed as previously described (Tong et al., 2012). Briefly, oocyte cumulus complexes were identified and washed in G-IVF (Vitrolife, Sweden), then inseminated with 50,000—100,000 normal motile spermatozoa, oocytes were examined for successful fertilization after 16-18h, zygotes with two pronuclei were allowed to continue culture for 48 h in G1 PLUS medium. On day 3, embryos were observed for morphology and blastomere numbers, embryos with 2-5 cells at day 3 were considered as possible arrested embryos. Arrested embryos were cultured in G2 PLUS medium for a further day (Vitrolife, Sweden), and if there was no further development then the embryos were defined as developmentally arrested. Arrested embryos were left untreated or treated with resveratrol (MCE, USA) or nicotinamide riboside/NR (Sigma-Aldrich, USA), in the resveratrol group, embryos were cultured with 1 *µ*M resveratrol for 6h, and then transferred into fresh G2 PLUS medium for 42 h. In the NR-treated embryos, embryos were cultured with 1 mM NR for 24 h, and then transferred into fresh G2 PLUS for 24 h, in the control group, embryos were cultured with fresh G2 PLUS for 48 h. Embryos were also treated with 0.5 *µ*M rapamycin (Sigma-Aldrich), 0.5 *µ*M PD0325901 (Selleck), and 25 *µ*g/ml vitamin C (Sigma-Aldrich), for 24 h, then transferred into fresh G2 PLUS medium for a further 24 h. The morphology and blastomere numbers were observed on day 4 and day 5. Embryos were culture in a tri-gas incubator with an environment of 6% CO_2_, 5% O_2_ and 89% N_2_ at 37°C.

### Immunofluorescence

After culture, embryos were fixed with 4% (w/v) paraformaldehyde for 30 minutes at room temperature, followed by washing in PBS (Phosphate-buffered saline) for three times. Embryos were permeabilized in 0.5% Triton X-100 for 30 minutes at room temperature, and blocked with blocking buffer (Beyotime, China) for 1 h at 37°C. Then embryos were then incubated with anti-SIRT1 antibody (1:100, Abcam, #ab189494, USA), anti-NRF2 antibody (1:100, Abcam, #ab137550, USA), anti-phospho-AKT (Ser473) antibody (1:200, CST, #4060S, USA), anti-phospho-P53 (Ser15) antibody (1:400, CST, #82530s, USA) or anti-p21 antibody (1:100, Abcam, #ab109520 USA) at 4°C overnight. Embryos were subsequently incubated with AlexaFluor-488 secondary antibody (1:500, Abcam, # ab150077, USA) for 2 h at 37°C, then washed three times. The DNA was stained with 4,6-diamino-2-phenyl indole (DAPI). Finally, the embryos were suspended in a microdrop with blocking solution and photographed with a fluorescence microscope (MetaSystems, Germany).

### Single embryo RNA-seq

Single embryo RNA-seq was performed essentially as described (Picelli et al., 2014). Briefly, single embryos were placed into 10 *µ*l of SMART-seq2 lysis buffer (1 ul RNAse inhibitor, 0.2% (v/v) Triton X-100) and stored at −80°C before sequencing. Embryos were sequenced on an Illumina sequencer according to the manufacturer’s instructions.

### RNA-seq data analysis

RNA-seq data was analyzed essentially as described in (Hutchins et al., 2017), with the exception that the STAR aligner was used (Dobin et al., 2013), RSEM was replaced with scTE (He et al., 2021), and GENCODE v32 (Frankish et al., 2021) was used for the transcript annotations. Data was GC-content normalized with EDASeq (Risso et al., 2011). Differential gene expression was determined using DESeq2 (Love et al., 2014), genes were defined as significantly differently expressed (DE) if they changed by at least 4-fold, and had a q-value (Benjamini & Hochberg corrected p-value) of 0.01 or less. GSEA was performed using fgsea, and all of the GSEAs shown in the manuscript had a q-value (Bonferroni-Hochberg corrected p-value, from 10,000 permutations) of less than 0.05 (Korotkevich et al., 2021). CytoTRACE (Gulati et al., 2020), was performed using default parameters on the unnormalized raw tag count matrix. Karyotype was estimated from the RNA-seq data using the software from https://github.com/MarioniLab/Aneuploidy2017 (Griffiths et al., 2017). The presumed normal samples are from a reanalysis of GSE66507 (Blakeley et al., 2015), PRJEB11202 (Petropoulos et al., 2016) and GSE36552 (Yan et al., 2013) embryo RNA-seq data. Other analysis was performed using glbase3 (Hutchins et al., 2014).

## Data Availability

The datasets supporting the conclusions of this article are available in the GSA (Genome Sequence Archive): HRA001406 under controlled access for human samples. The authors declare that the data supporting the findings of this study are available within the article and its Supplementary Information files, or from the corresponding author upon reasonable request.

## Statistics

Statistical analysis was used for the differential expression measures. Differential expression was determined using DESeq2 (Love et al., 2014), and minimum thresholds of >4-fold change, and a q-value of 0.01 (Benjamini-Hochberg corrected p-value) was used to determine significantly different. For GSEA, a minimum q-value of 0.05 (Benjamini-Hochberg corrected p-value) was considered significant, as estimated by random permutation by fgsea (Korotkevich et al., 2021).

## Study Approval

This study was approved by Ethical Approval Board (2020-866-75-01) of the Shuguang Hospital affiliated to Shanghai University of Traditional Chinese Medicine, and Southern University of Science and Technology ethical committee (2021SWX012). Arrested embryos used in this study were rejected embryos from normal IVF procedures, and were used with the patients informed, written consent. No normal control embryos (non-arrested) were consumed in this study.

## Acknowledgements

This work was supported by the National Key R&D Program of China (2018YFC1704300), the National Natural Science Foundation of China (81070494, 81170571, 81571442 and 31970589), the Shenzhen Innovation Committee of Science and Technology (JCYJ20200109141018712 and ZDSYS20200811144002008 to the Shenzhen Key Laboratory of Gene Regulation and Systems Biology), and the Stable Support Plan Program of the Shenzhen Natural Science Fund (20200925153035002).

## Author contributions

Y.Y. Y.L., P.Y., Z.W., performed the embryo manipulations and experiments on embryos, and Y.Y. led this effort. S.L., Z.Y., I.A.B., and A.P.H. performed the bioinformatic analysis, and S.L. led this effort. M.G., X.F., L.Y., Z.Y., S.L., performed sequencing quality control. All other authors assisted with manuscript preparation. G.T conceived of the project. A.P.H. and G.T. funded the study and wrote the manuscript with assistance from all authors.

