## Supplementary Figures for "Human embryos arrest in a quiescent-like state characterized by metabolic and zygotic genome activation problems"

#### Supplementary Material

**Supplementary Table 1** – Differentially expressed genes defined as major ZGA genes (up-regulated), and maternal RNA-clearance genes (down-regulated) (From **Supplementary Fig. 3a**).

**Supplementary Table 2** – Differentially expressed genes (fold-change > 4, q-value <0.01) for all Type I, II and III arrested embryos.

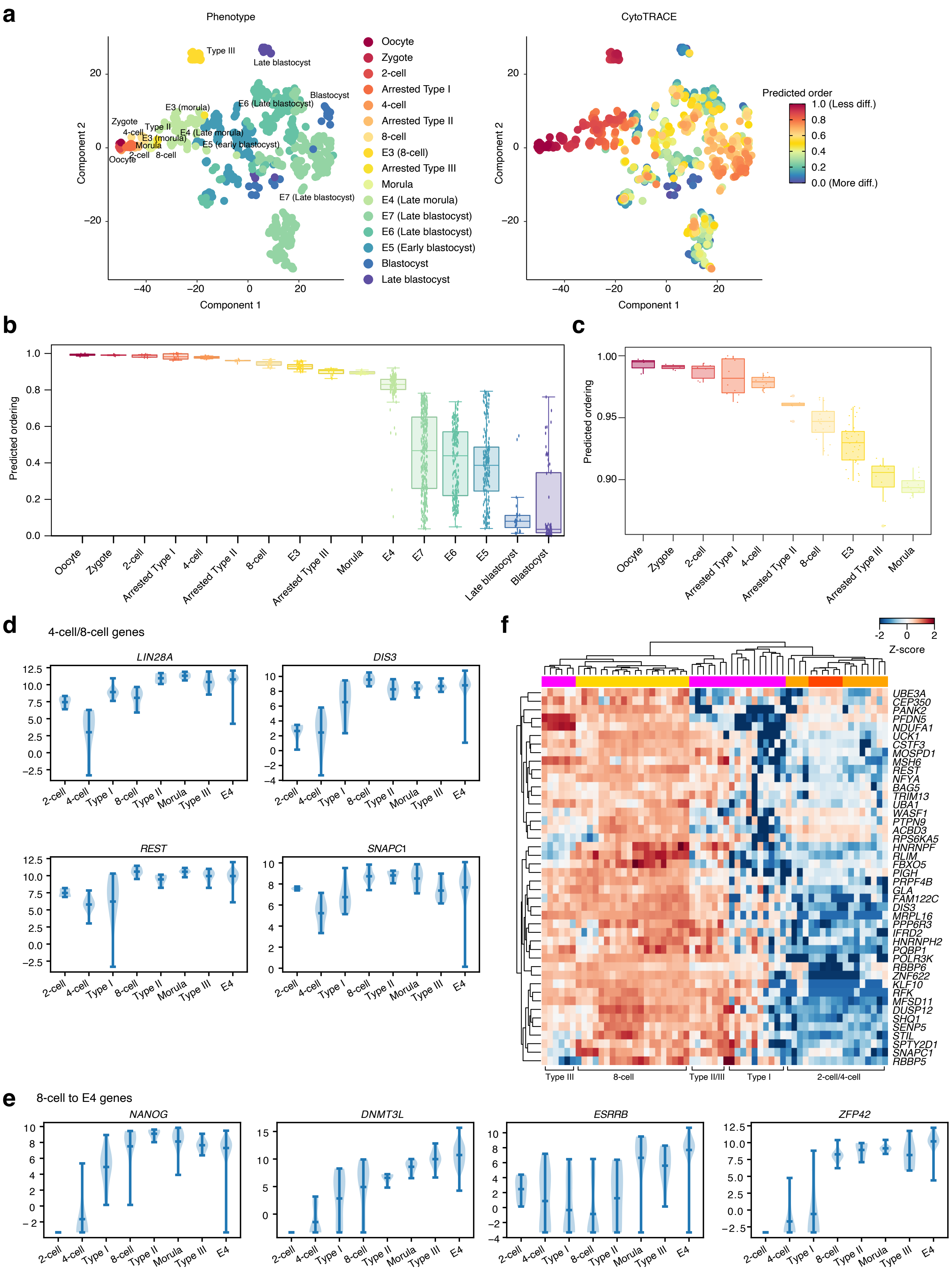


**Supplementary Figure 1**

**Supplementary Figure 1**. Arrested embryos maintain developmental potential

**(A)** CytoTRACE cell embedding manifold, colored by cell type and embryo-stage (left) or by predicted developmental order (right). A predicted developmental order score of 1.0 is less differentiated, and a score of 0.0 is more. E = Embryonic-stage samples, as defined in (Petropoulos et al., 2016), for this and all subsequent figures.

**(B)** Box plots of the predicted ordering from CytoTRACE for all cells/embryos at the indicated stages, ordered by the mean developmental predicted ordering. Each dot is a cell/embryo.

**(C)** As in **panel B**, but only showing the stages from oocyte to morula.

**(D)** Violin plots showing expression of 4-cell/8-cell-specific human genes.

**(E)** Violin plots showing expression of 8-cell/E4-specific human genes, i.e. developmental genes involved in the establishment of the blastocyst.

**(F)** Heatmap of the Z-scores of expression of the genes in the 8-cell-signature identified in (Hasegawa et al., 2015).

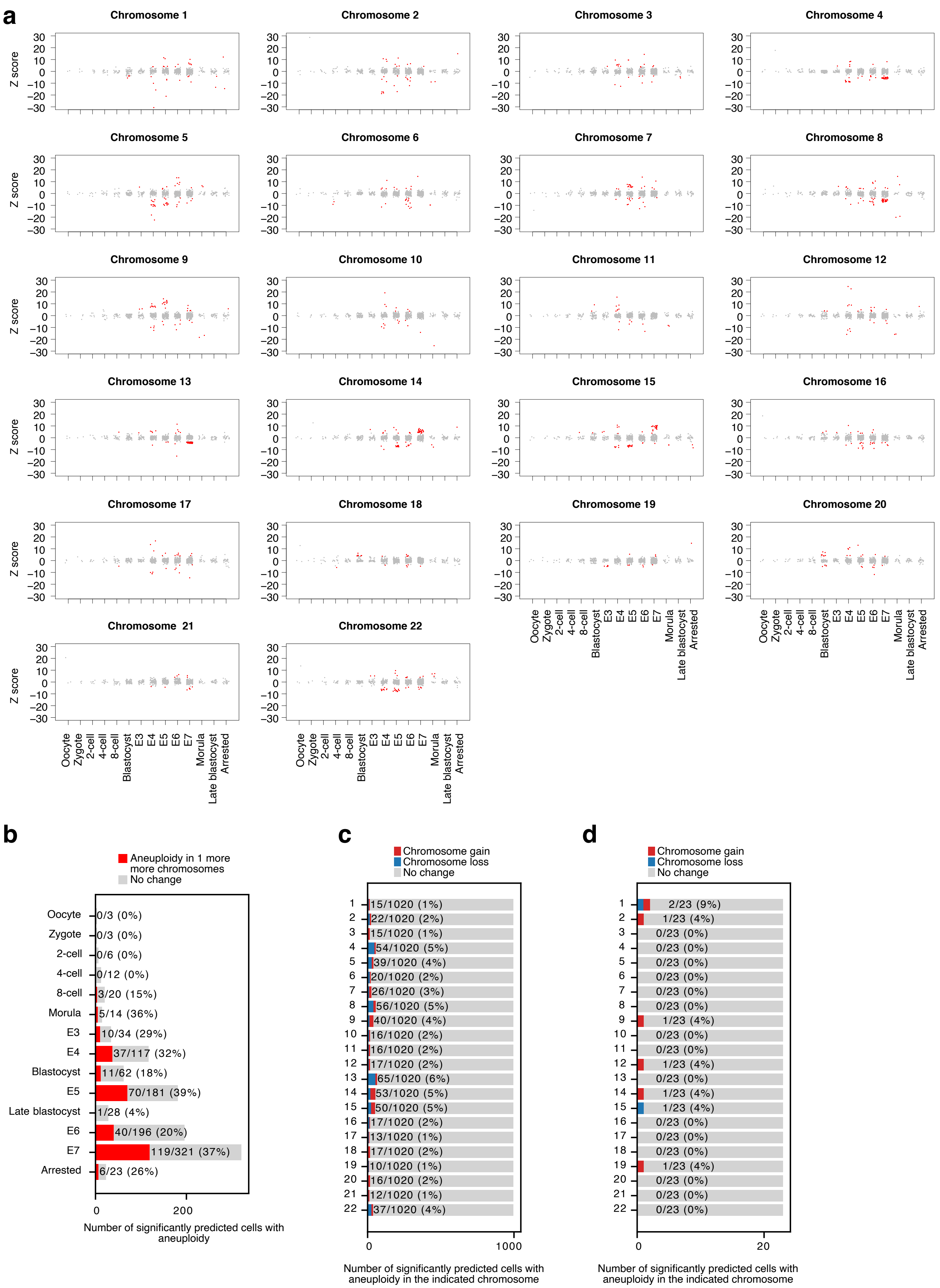


### Supplementary Figure 2

**Supplementary Figure 2**. Arrested embryos are (mainly) karyotypically normal

**(A)** Karyotype abnormalities estimated using the approach outlined in (Griffiths et al., 2017). In these plots, each chromosome is plotted separately, and the grey dots are each cell/embryo in the indicated stage of development or arrest. Red dots indicate when the gene expression on that chromosome exceeds the Z-score threshold and is significantly over or under represented. Red dots are indicative of aneuploidies, and those above the line suggest a gain of a chromosome or part of a chromosome, and those below the line suggest a loss of a chromosome or part.

**(B)** Bar chart showing the predicted aneuploidies in the indicated developmental stages. Red bars indicate cells/embryos with a predicted loss or gain of a chromosome, whilst those in grey are predicted to be normal.

**(C)** Percentage of the aneuploidies observed, broken down by chromosome in the normal embryo data set.

**(D)** As in **panel C**, but only the arrested embryos.

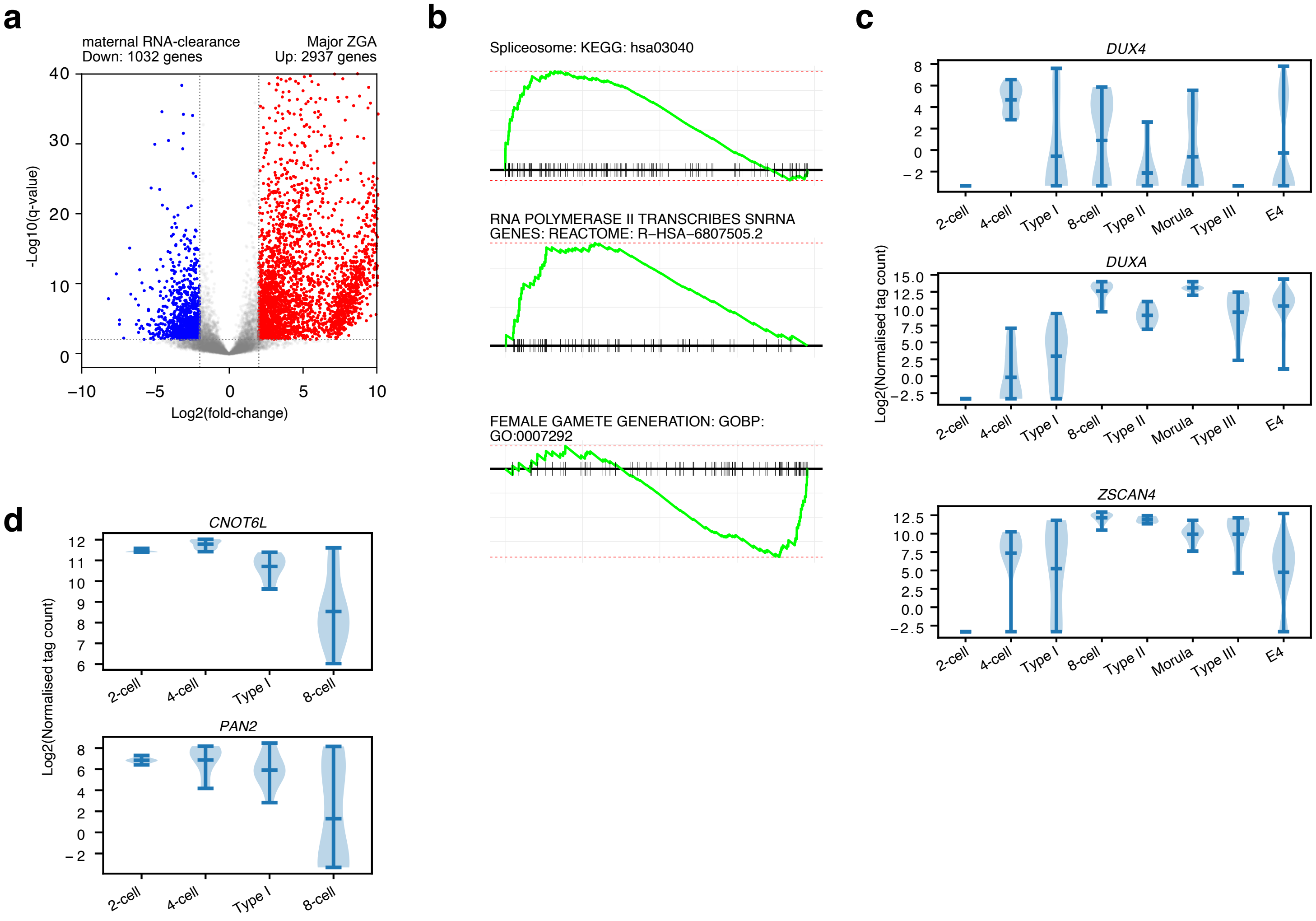


### Supplementary Figure 3

**Supplementary Figure 3**. Type I arrested embryos have MZT problems.

- **(A)** Volcano plot showing the fold-change versus significance when comparing 2-cell-stage embryos versus 8-cell-stage. Differential expression was calculated using DESeq2, and a minimum fold-change of 4, and a q-value of 0.01 was considered significantly different. The q-value is the Bonferroni-Hochberg multiple test corrected p-value. We defined 'maternal RNA-clearance genes' as those that were significantly down-regulated, and 'major ZGA genes' as those that were up-regulated. Significantly differentially expressed up-regulated genes are labelled in red, and down-regulated in blue. The number of genes passing the differential expression thresholds are marked on the plot.
- **(B)** GSEA for the up and down-regulated genes as ranked in **panel A**.
- **(C)** Violin plots for the expression of key major ZGA genes, *DUX4*, *DUXA* and *ZSCAN4*.
- **(D)** Violin plots for the expression of critical maternal RNA clearance genes *CNOT6L* and *PAN2*.

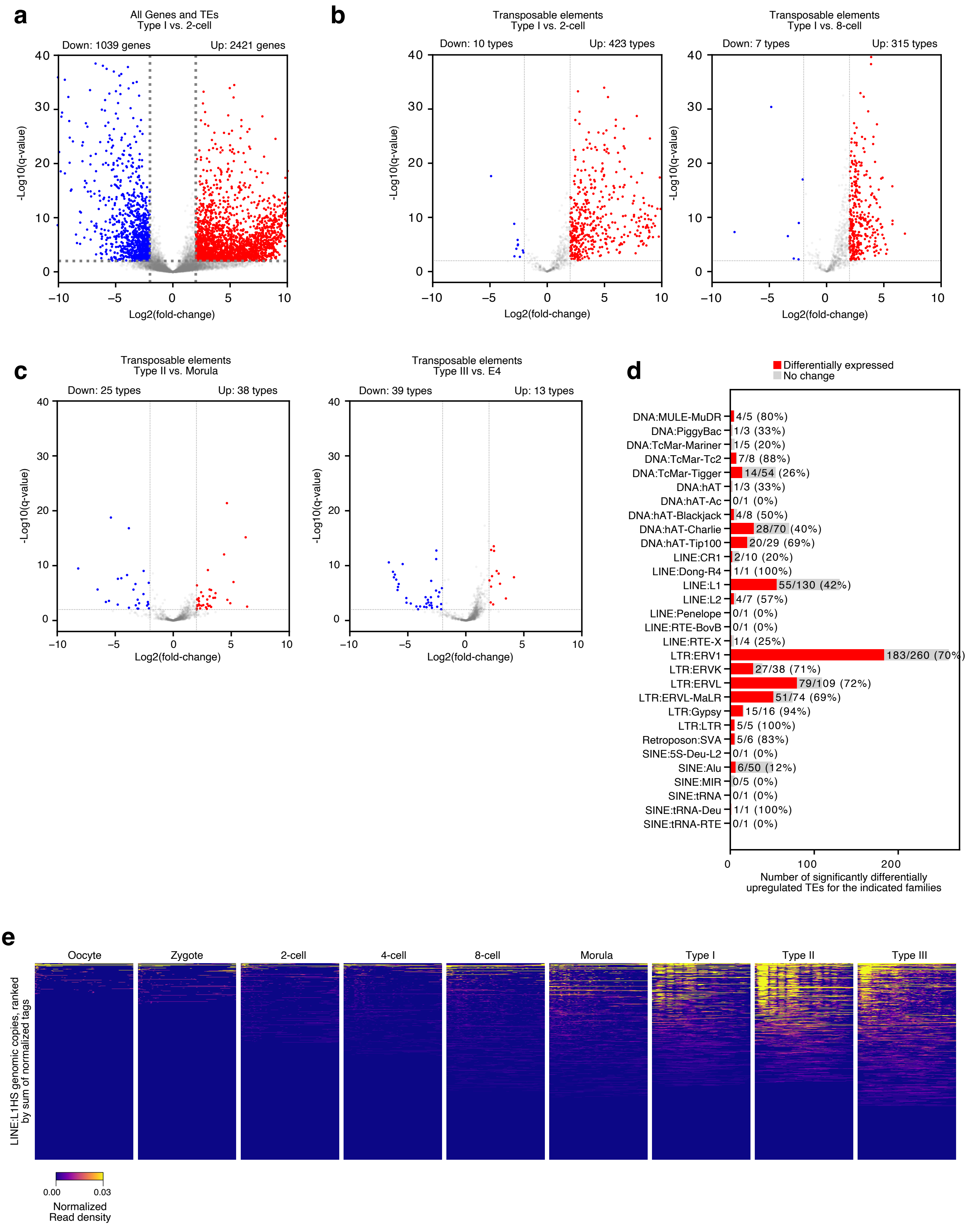


### Supplementary Figure 4

**Supplementary Figure 4**. Transposable element expression is disturbed in Type I arrested embryos.

- **(A)** Volcano plot for all genes and transposable elements (TEs) when comparing Type I arrested embryos to 2-cell-stage cells. Differential expression was calculated using DESeq2, and a minimum fold-change of 4, and a q-value of 0.01 was considered significantly different. The q-value is the Bonferroni-Hochberg multiple test corrected p-value. Significantly differentially expressed up-regulated genes are labelled in red, and down-regulated in blue. The number of genes passing the differential expression thresholds are marked on the plot.
- **(B)** Volcano plot, as in **panel A**, but only containing TE types, and comparing Type I arrested embryos to 2-cell-stage embryos (left volcano), or 8-cell-stage embryos (right volcano).
- **(C)** Volcano plot, as in **panel A**, but only containing TE types, and comparing Type II arrested embryos to morula-stage embryos (left volcano), or Type III arrested embryos versus E4 (early blastocyst)-stage embryos (right volcano).
- **(D)** Number of differentially regulated TE types for the indicated TEs, when comparing Type I arrested embryos to 4-cell-stage embryos.
- **(E)** Heatmaps of the RNA-seq read tag density for all genomic copies of LINE L1HS copies (rows). Heatmaps are the density of normalized tag counts (in reads per million) for each sample, and are ranked by the sum of each row for each heatmap.

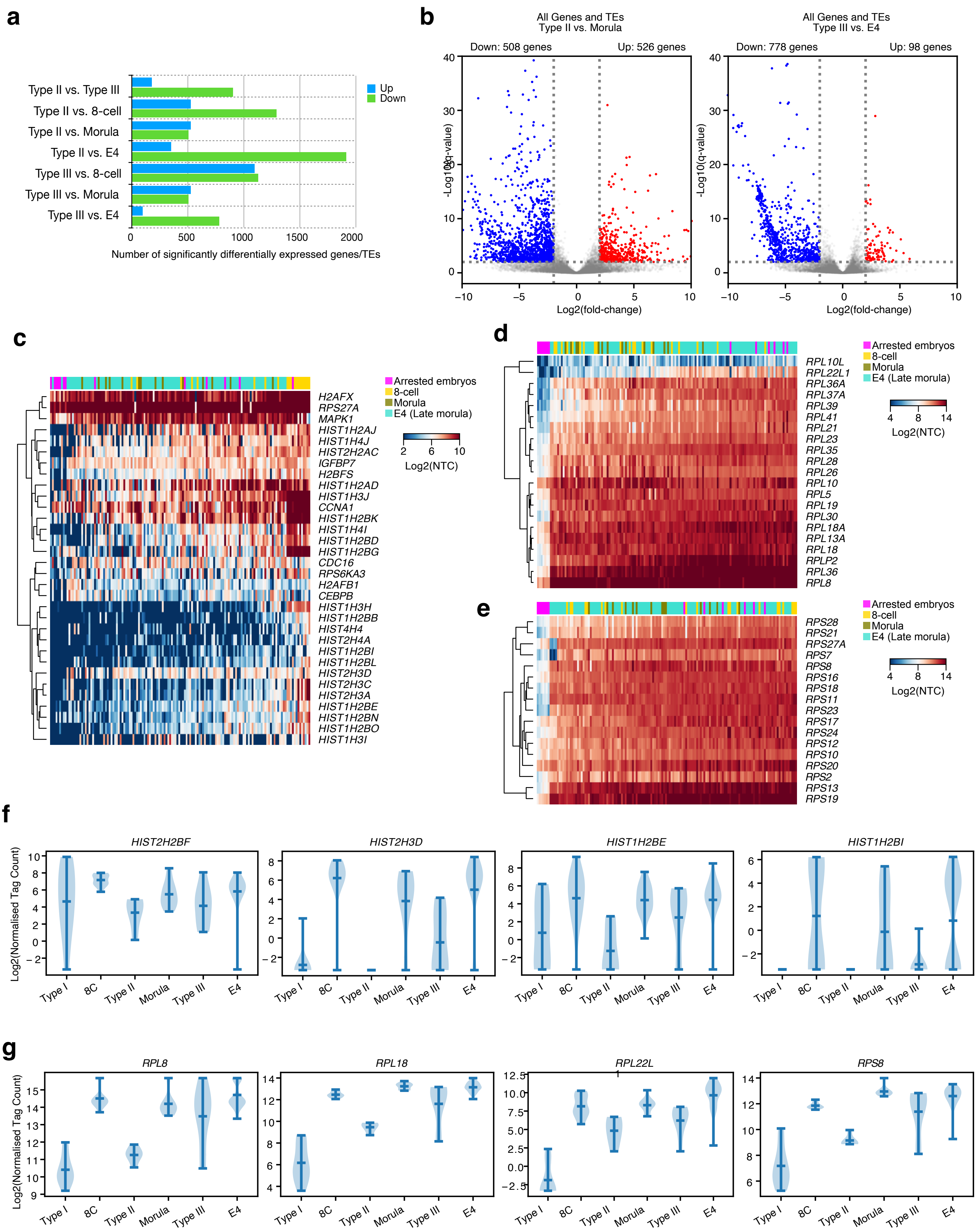


### Supplementary Figure 5

**Supplementary Figure 5**. Arrested embryos have reduced ribosome and nucleosome expression.

- **(A)** Number of significantly differentially regulated genes and TEs (A fold-change of at least 4, and a Bonferroni-Hochberg corrected q-value of less than 0.01) in the indicated comparisons.
- **(A)** Volcano plots showing all genes and TEs when comparing Type II versus morula (left) or Type III versus E4 (right) -stage embryos. Differential expression was calculated using DESeq2, and a minimum fold-change of 4, and a q-value of 0.01 was considered significantly different. The q-value is the Bonferroni-Hochberg multiple test corrected p-value. Significantly differentially expressed up-regulated genes are labelled in red, and down-regulated in blue. The number of genes passing the differential expression thresholds are marked on the plot.
- **(B)** Heatmap of the expression of all histones/nucleosomes and selected senescence-related genes (From the REACTOME category: SENESCENCE−ASSOCIATED SECRETORY PHENOTYPE (SASP): REACTOME: R−HSA−2559582.2). Expression is presented as log2 normalized tag count (NTC).
- **(C)** Heatmap of all significantly differentially expressed (fold-change >4 and q-value <0.01) large ribosome subunits. Expression is presented as log2 normalized tag count (NTC).
- **(D)** As in **panel D**, but for all significantly differentially expressed small ribosome subunits.
- **(E)** Violin plot for the expression of selected histone proteins.
- **(F)** Violin plot for selected large or small ribosome subunits.

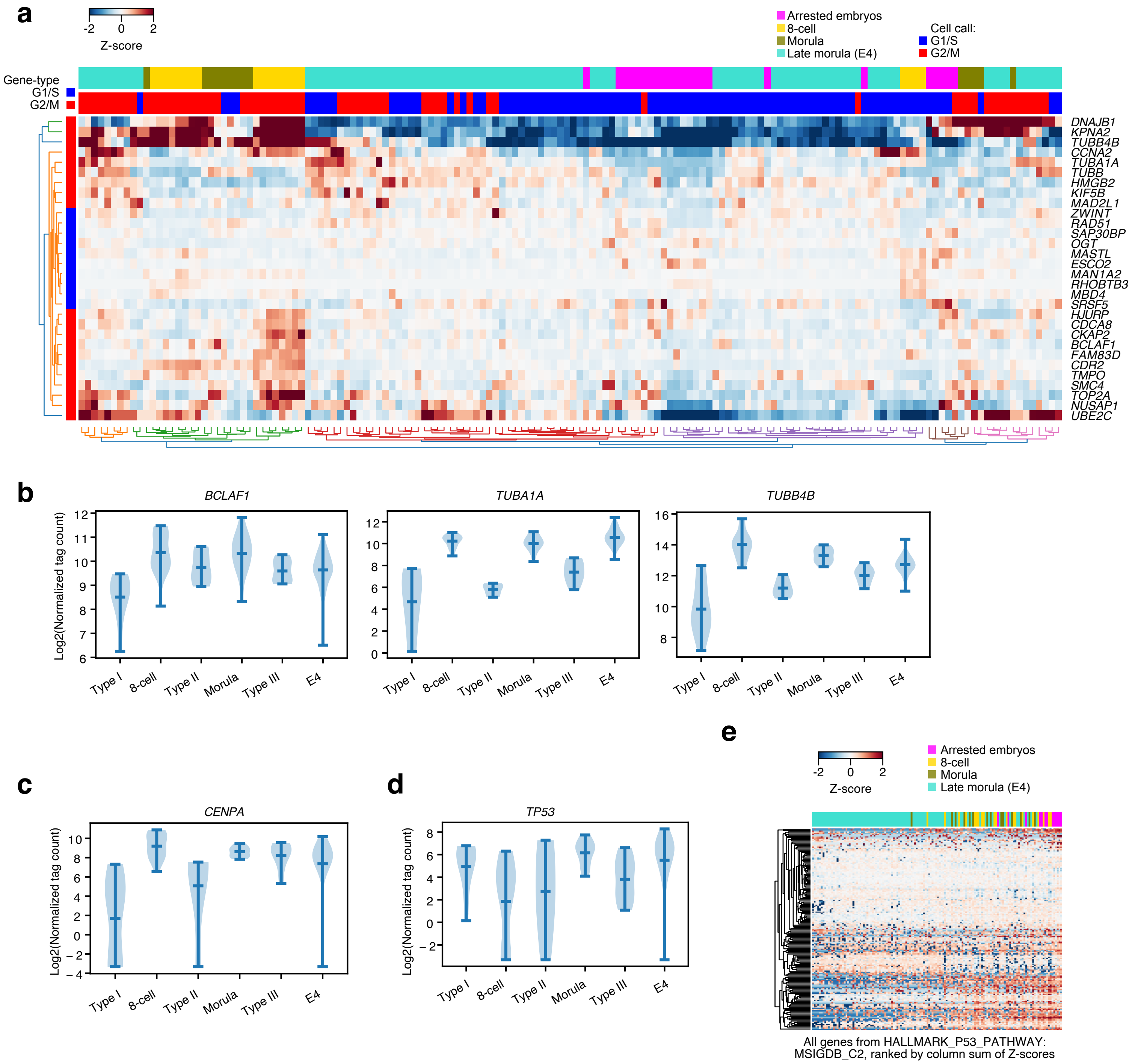


**Supplementary Figure 6**

**Supplementary Figure 6**. Arrested embryos have decreased cell cycle activity.

- **(A)** Heatmap showing the expression of selected cell cycle-related genes. The cell/embryo stage is indicated in the top, colored bar legend. The predicted cell cycle phase is in red and blue, below.
- **(B)** Violin plots showing expression of cell cycle-related gene, *BCLAF1*, and the tubulin subunits *TUBA1A* and *TUBB4B*.
- **(C)** Violin plot for the expression of *CENPA*.
- **(D)** Violin plot for the expression of *TP53* (p53).
- **(E)** Heatmap for the expression of p53 target genes (HALLMARK_P53_PATHWAY set), ranked by the sum of the columns. Each column is a single cell or embryo, and the arrested embryos are labelled in pink.

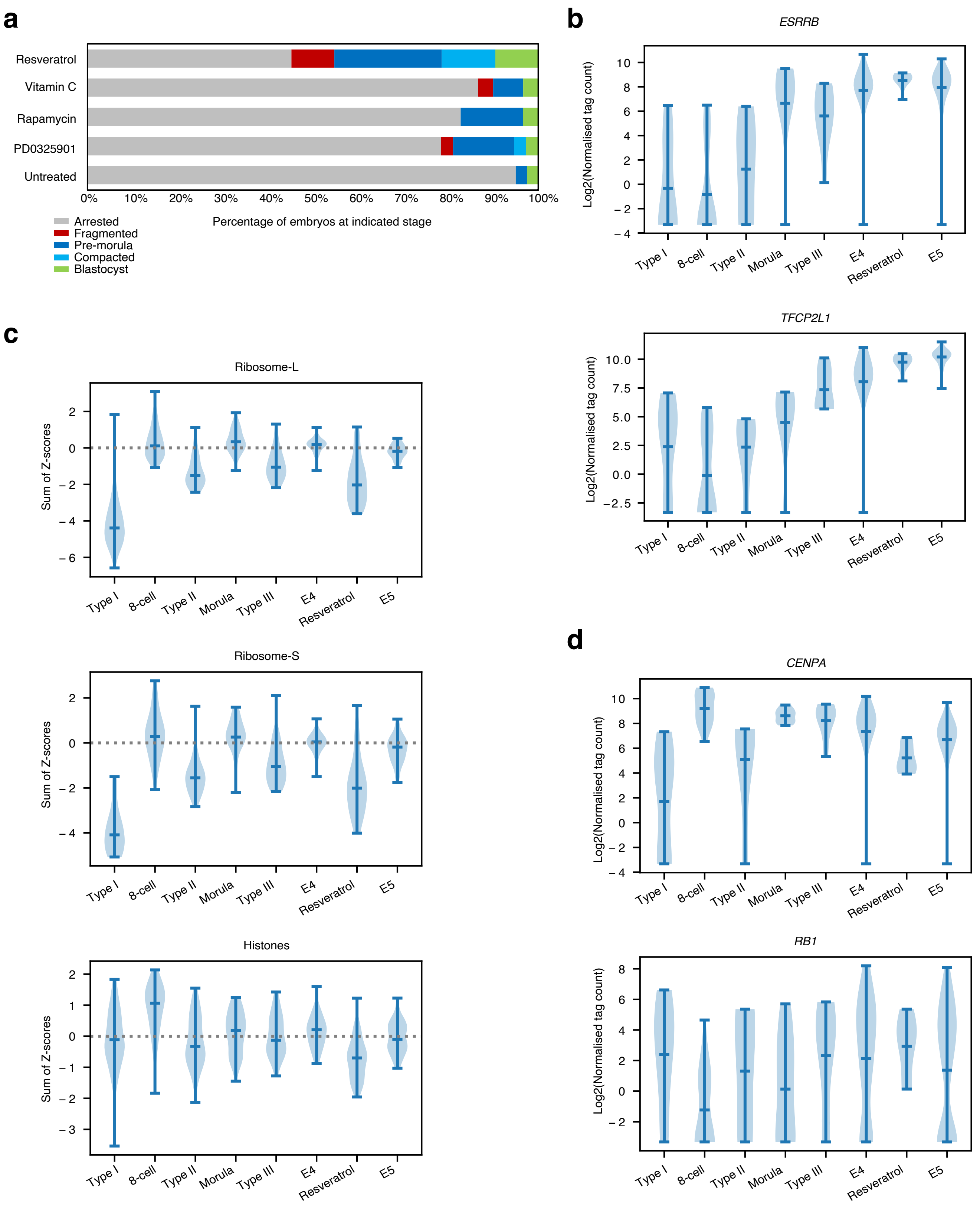


### Supplementary Figure 7

**Supplementary Figure 7**. Treatment of arrested embryos with small molecules; and resveratrol corrects developmental and cell cycle problems, but not ribosomes and nucleosome expression.

- **(A)** Percentage of arrested embryos that recommenced development when treated with the indicated small molecules.
- **(B)** Violin plot showing the expression of the blastocyst-related genes *ESRRB* and *TFCP2L1* in the indicated embryonic stages and in arrested and resveratrol-treated embryos.
- **(C)** Violin plots showing the distribution of Z-scores of expression for all the small and large ribosomes and histone genes in the indicated embryonic stages and in arrested and resveratrol-treated embryos.
- **(D)** Violin plots showing the expression of the cell cycle-related genes *CENPA* and *RB1* in the indicated embryonic stages and in arrested and resveratrol-treated embryos.

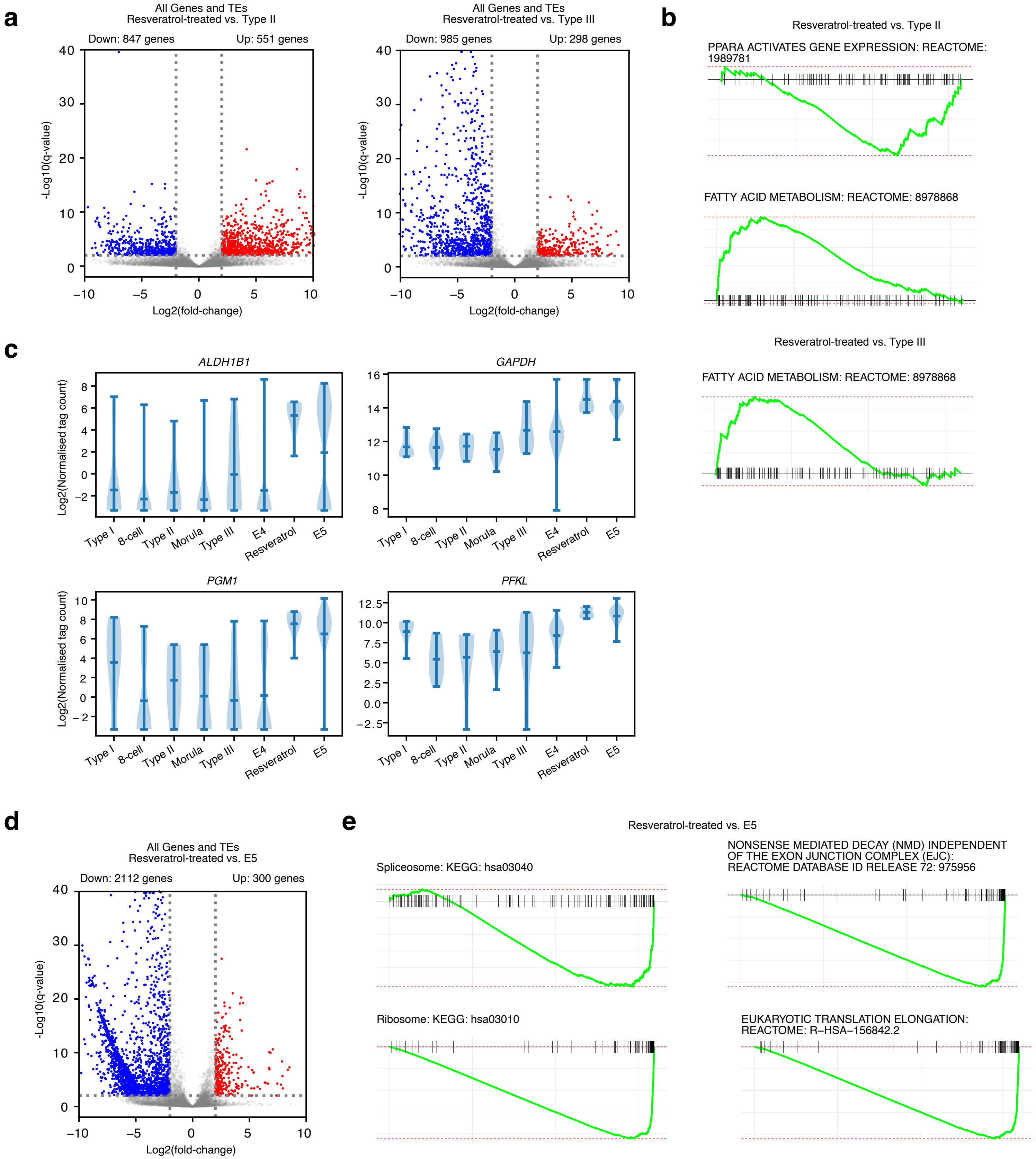


**Supplementary Figure 8**

**Supplementary Figure 8**. Comparison of resveratrol-treated versus other embryonic cells.

- **(A)** Volcano plot showing all significantly differentially expressed (fold-change >4 and q-value <0.01) for all genes and TEs when comparing showing resveratrol versus Type II (left) or Type III (right)-arrested embryos. Differential expression was calculated using DESeq2, and a minimum fold-change of 4, and a q-value of 0.01 was considered significantly different. The q-value is the Bonferroni-Hochberg multiple test corrected p-value. Significantly differentially expressed up-regulated genes are labelled in red, and down-regulated in blue. The number of genes passing the differential expression thresholds are marked on the plot.
- **(B)** GSEA showing significantly different terms for resveratrol versus Type II arrested embryos.
- **(C)** Violin plots showing the expression of the indicated glycolysis-related genes *ALDH1B1*, *GAPDH*, *PGM1* and *PFKL*.
- **(D)** Volcano plots (as in **panel A**), but showing Resveratrol-treated embryos versus E5-stage embryos.
- **(E)** GSEA of the ranked differentially expressed genes from **panel D**.

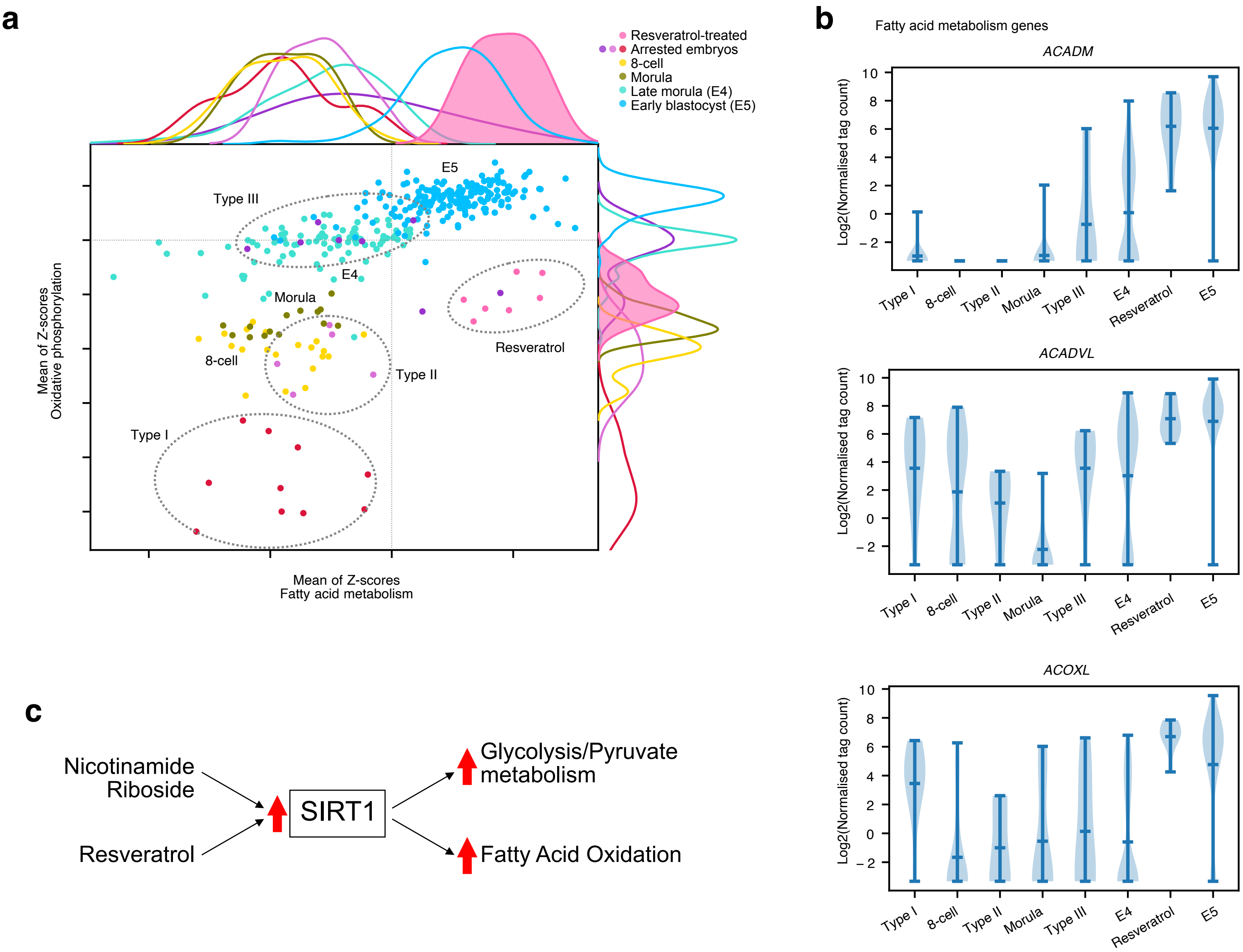


#### Supplementary Figure 9

**Supplementary Figure 9**. Resveratrol upregulates the expression of fatty acid metabolic genes.

- **(A)** 2D dotplot showing the sum of the Z-scores for the genes in the indicated KEGG categories. The x-axis scores the fatty acid metabolism pathway, the y-axis the oxidative phosphorylation pathway. Each dot in the plot is a cell/embryo, and the top and right axis have the kernel density for each group of cells. The resveratrol-treated embryos have a filled in color (pink) for emphasis. The arrested and resveratrol-treated embryos are indicated by dashed lines, and the normal developmental states are indicated by labels.
- **(B)** Violin plots showing the expression of selected fatty acid-related metabolic genes.
- **(C)** A model for the action of resveratrol and NR on SIRTs and metabolic pathways.

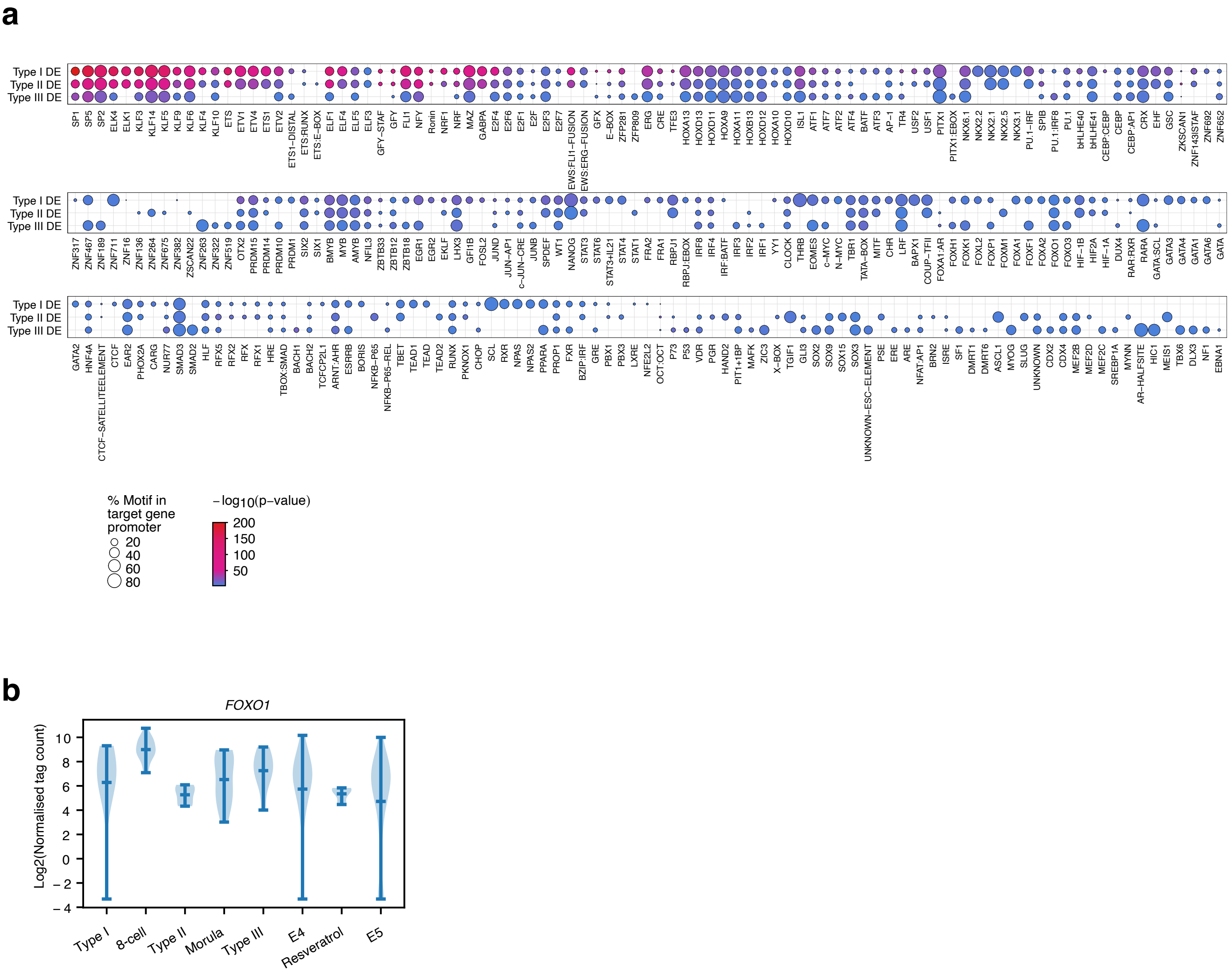


### Supplementary Figure 10

**Supplementary Figure 10**. Transcriptional regulation of embryonic arrest.

- **(A)** All significantly enriched motifs in the Type I, II, III-arrested embryos in the promoters of differentially expressed (DE) genes. Motif discovery was performed using HOMER with default settings (Heinz et al., 2010), against the promoters (defined here as -1000 bp upstream) of the DE genes in the indicated types of arrested-embryo. The size of the circle indicates the percent of gene promoters that had the motif, and the color indicates the p-value for enrichment.
- **(B)** Violin plot showing the expression of *FOXO1* in the indicated embryonic cell types.
